# Essential role for FtsL in activation of septal PG synthesis

**DOI:** 10.1101/2020.09.01.275982

**Authors:** Kyung-Tae Park, Shishen Du, Joe Lutkenhaus

## Abstract

Spatiotemporal regulation of septal PG synthesis is achieved by coupling assembly and activation of the synthetic enzymes (FtsWI) to the Z ring, a cytoskeletal element required for division in most bacteria. In *E. coli* the recruitment of the FtsWI complex is dependent upon the cytoplasmic domain of FtsL, a component of the conserved FtsQLB complex. Once assembled, FtsWI is activated by the arrival of FtsN, which acts through FtsQLB and FtsA that are also essential for their recruitment. However, the mechanism of activation of FtsWI by FtsN is not clear. Here, we identify a region of FtsL that plays a key role in the activation of FtsWI which we designate AWI (<u>A</u>ctivation of Fts<u>WI</u>) and present evidence that FtsL acts through FtsI. Our results suggest that FtsN switches FtsQLB from a recruitment complex to an activator with FtsL interacting with FtsI to activate FtsW. Since FtsQLB and FtsWI are widely conserved in bacteria this mechanism is likely to be also widely conserved.

**Significance:** A critical step in bacterial cytokinesis is the activation of septal peptidoglycan synthesis at the Z ring. Although FtsN is the trigger and acts through FtsQLB and FtsA to activate FtsWI the mechanism is unclear. Here we find an essential role for FtsL in activating septal PG synthesis and find that it acts on FtsI. Our results suggest a model where FtsWI is recruited in an inactive form by FtsQLB and upon FtsN arrival, FtsQLB undergoes a conformational change so that a region of FtsL, that we designate the AWI domain, becomes available to interact with FtsI and activate the FtsWI complex. This mechanism for activation of the divisome has similarities to activation of the elongasome and is likely to be widely conserved in bacteria.

## Introduction

Bacterial cell division in most bacteria is carried out by a large protein complex called the divisome or septal ring (1, 2). In *E. coli* it consists of 12 essential proteins and an ever-increasing number of nonessential proteins. The essential proteins include FtsZ, which assembles into treadmilling filaments that are tethered to the membrane by FtsA and ZipA (Z ring), and 9 additional proteins which display the following dependency for recruitment – FtsE/X < FtsK < FtsQ < FtsL/B < FtsW < FtsI and FtsN (Fig. 1) (1-4). Among these, FtsW is a newly described glycosyltransferase of the SEDS family that works in concert with a transpeptidase (FtsI [PBP3]) to synthesize septal PG (5-8). A key step in cell division is the activation of these enzymes by the last arriving essential protein FtsN (3, 9) which acts through FtsA and the FtsQLB complex (10-12).

**Fig. 1.**
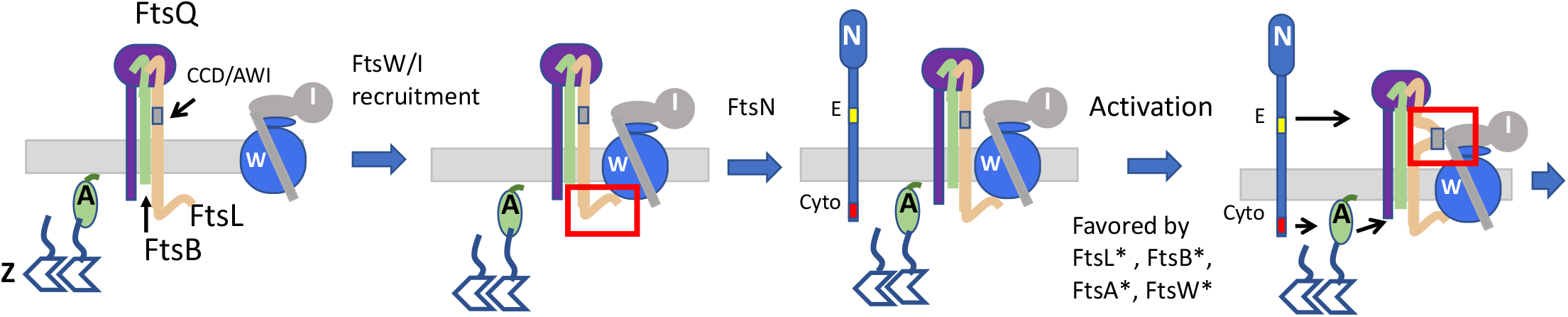
Assembly and activation of septal PG synthesis. Model for recruitment and activation of FtsWI. FtsQLB localizes to the Z ring and recruits FtsW in a ^cyto^FtsL dependent manner with FtsW recruiting FtsI. FtsN arrives and ^cyto^FtsN interacts with FtsA in the cytoplasm and the ^E^FtsN domain acts in the periplasm. Together they switch FtsQLB from the OFF to the ON conformation resulting in activation of FtsWI. In the model activation occurs when the AWI (activation of FtsWI) domain (which overlaps the CCD [control of cell division] in FtsL) within the FtsQLB complex becomes available to contact FtsI. Activation mutations (*) in *ftsA, ftsB, ftsL* and *ftsW* require less FtsN. FtsB* and FtsL* are thought to switch FtsQLB to the ON state. FtsA* may also do this whereas FtsW* is likely to lead to an enzymatically active form of FtsW. For simplicity ZipA, FtsEX and FtsK are not depicted. FtsZ, FtsA, FtsW, FtsI and FtsN are indicated by single letters. The red rectangles highlight interactions between FtsL and FtsWI. FtsZ, FtsA, FtsW, FtsI and FtsN are indicated by single letters.

The FtsQLB complex is widely conserved among peptidoglycan containing bacteria and links the Z ring to the septal PG synthesis machinery (FtsWI) (13). Each protein in the FtsQLB complex is a bitopic membrane protein with a short cytoplasmic region connected to a larger periplasmic domain by a single transmembrane domain. FtsQ targets the FtsQLB complex to the Z ring in an FtsK dependent fashion and the cytoplasmic domain of FtsL is required to recruit FtsW(3) (Fig. 1). FtsL and FtsB form a multimer with interactions occurring between their alpha-helical transmembrane domains as well as their putative periplasmic coiled-coil domains (14-17). They also interact with FtsQ through their C-terminal domains that lie beyond the coiled-coil domains forming a 1:1:1 complex which may dimerize (13, 15, 18). Recently, a peptide corresponding to the C-terminal region of FtsB was crystallized bound to FtsQ (19, 20).

Activation of FtsWI by FtsN requires two domains of FtsN; the ^cyto^FtsN domain acts on FtsA and the ^E^FtsN domain, a short putative helical segment in the periplasm, likely acts on FtsQLB (10, 21, 22) (Fig. 1). In a proposed model FtsN switches both FtsA and FtsQLB to an ON state which activates FtsWI (10, 11). This regulatory model is based in part upon the isolation of “activation (superfission)” mutations (requiring less FtsN) in *ftsL* and *ftsB* which identified a short periplasmic region in both proteins, designated CCD for control of cell division (10). The CCD connects the coiled-coil domain of each protein to its distal C-terminal region that binds to FtsQ (13, 18). Although it is not clear how these mutations work, it is likely they mimic FtsN action resulting in a change in conformation of the FtsQLB complex to the ON state that activates FtsWI (Fig. 1). Activation mutations have also been isolated in *ftsA* and *ftsW* (10, 12). Such mutations in *ftsA* could cause it to act on FtsQLB or FtsW whereas such mutations in *ftsW* could lead to an enzymatically active mutant. To address the mechanism of FtsWI activation we set out to isolate dominant negative mutations in *ftsL* and *ftsB*. Such mutations should yield a FtsQLB complex that no longer activates FtsWI and yield information about the activation mechanism. By exploring the effect of the dominant negative mutations, as well as the activation mutations, on the recruitment and activation of FtsWI we find an essential role for FtsL in the activation of FtsWI.

## Results

### Isolation of dominant negative mutations in *ftsL* but not *ftsB*

To isolate dominant negative mutations in *ftsL* and *ftsB*, they were subjected to random mutagenesis, cloned into plasmid downstream of an IPTG-inducible promoter and introduced into a wild type strain. Colonies were picked and screened for inhibition of growth by spotting on plates containing IPTG. Three strong dominant negative mutations were obtained in *ftsL* (*ftsL*^*E87K*^, *ftsL*^*L86F*^, and *ftsL*^*A90E*^) as well as two weak mutations (*ftsL*^*R61C*^ and *ftsL*^*L24K*^) but none were obtained in *fts*B (Fig. 2A and Table 1). Changing *ftsL*^*R61C*^ to *ftsL*^*R61E*^ resulted in a stronger dominant negative mutant (Table 1) while *ftsL*^*L24K*^ is discussed later. Induction of the *ftsL* alleles in liquid culture resulted in filamentation (Fig. 2B and Table 1). Complementation tests confirmed they were loss of function mutations as they were unable to complement a *ΔftsL* strain (Fig. S1A, Table 1). Interestingly, three of these mutations overlapped the CCD domain which was previously defined by activation mutations that decrease the dependency upon FtsN (10, 11) (Fig. 2C). Using site-directed mutagenesis we altered additional residues around the CCD and isolated three additional dominant negative mutations (*ftsL*^*R82E*^, *ftsL*^*N83K*^ and *ftsL*^*L84K*^) (Fig. 2C, Table 1). However, extending the mutagenesis to flanking regions as well as the C-terminal region of *ftsL* did not yield any additional dominant negative mutations (Fig. 2C and Table 1). Although the residues we identified overlap the CCD domain they are distinct from the residues involved in activation and are on opposite sides of a putative alpha helix. Since these mutations lead to a dominant negative effect, they behave as though they are nonresponsive to FtsN, the opposite of activation mutations (Fig. 2D). We designate the region identified by the dominant negative residues as AWI (<u>A</u>ctivation of Fts<u>WI</u>) based on results described below.

**Table 1.**
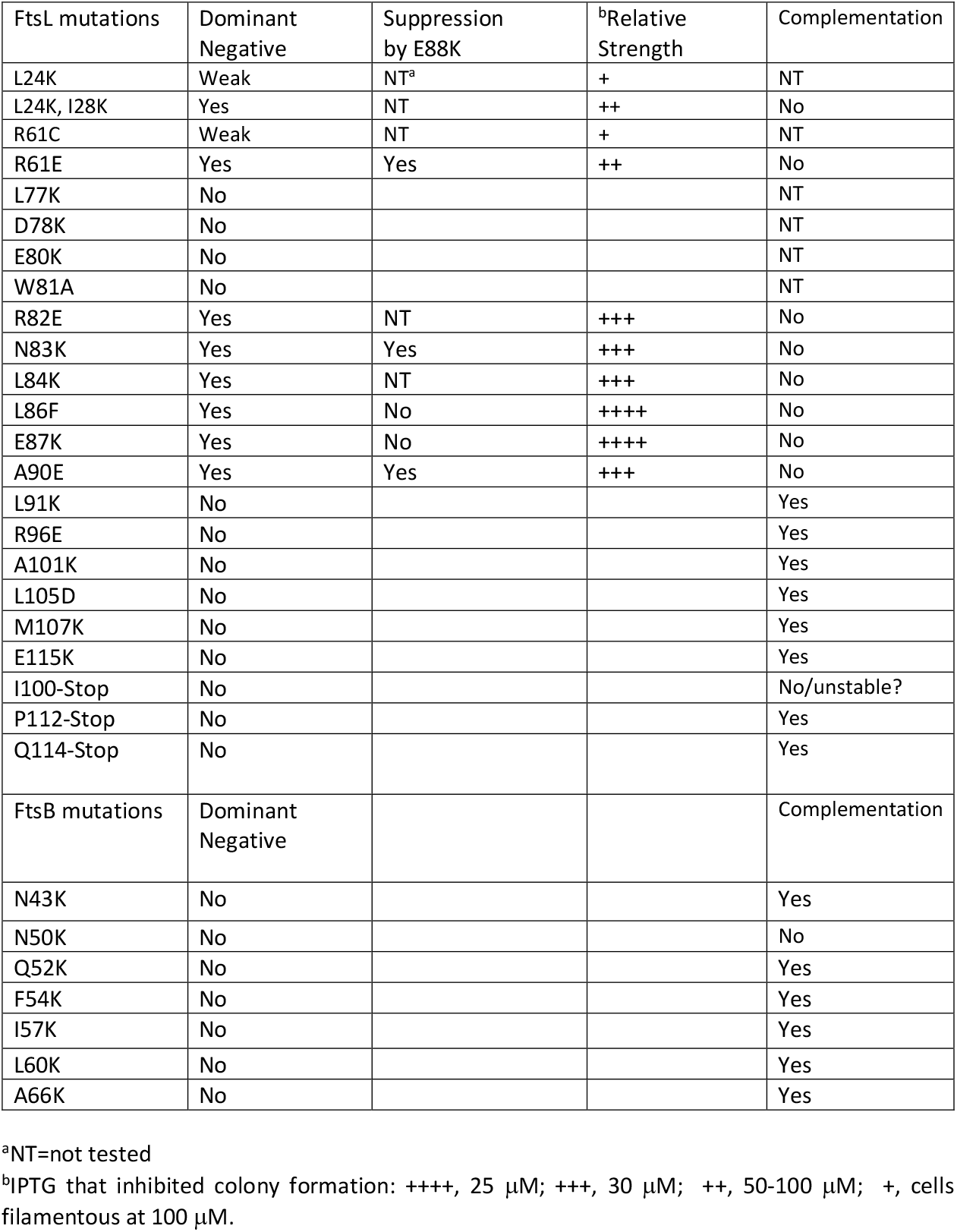
Summary of the point mutations in *ftsL* and *ftsB*

**Fig. 2.**
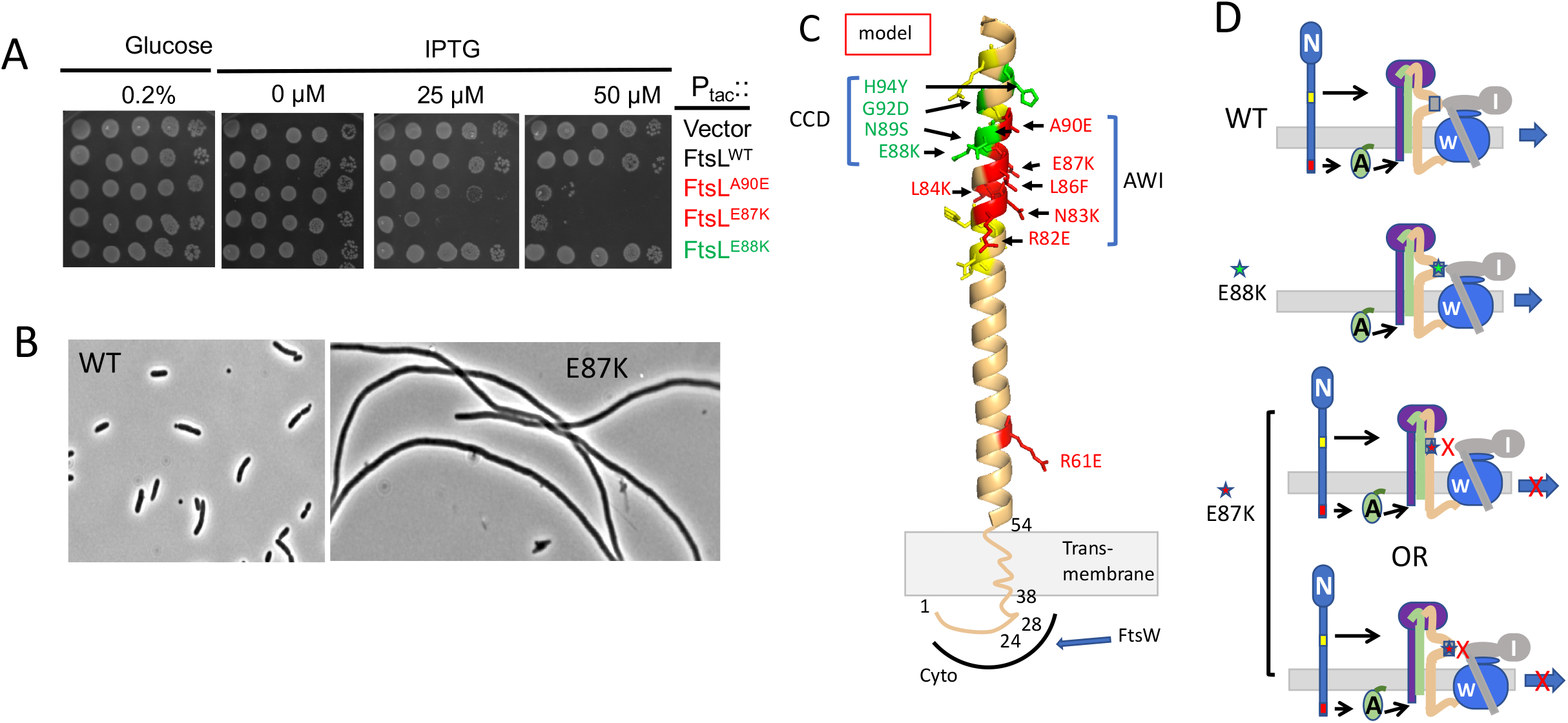
Isolation of dominant negative mutations in *ftsL*. A) Spot test of dominant negative mutations in *ftsL. ftsL* was subjected to random PCR mutagenesis, cloned downstream of the *tac* promoter in an expression vector containing an IPTG-inducible promoter (pJF118EH) and transformed into JS238. Colonies were picked and screened for sensitivity to IPTG. *ftsL*^*WT*^ and *ftsL*^*E88K*^ (an activation allele) were included as controls and are not toxic. Several strong dominant negative mutations (*ftsL*^*E87K*^, *ftsL*^*L86F*^ and *ftsL*^*A90E*^) and two weak mutations (*ftsL*^*R61C*^ and *ftsL*^*E24K*^) were obtained in this way (Table 1). Additional mutations were obtained by site-directed mutagenesis. B) Dominant negative mutants inhibit division. Phase contrast micrographs of JS238 expressing *ftsL* or *ftsL*^*E87K*^ (derivatives of pKTP100 [P_*tac*_::*ftsL*]) grown in liquid culture and induced with 50 µM IPTG for two hours. Induction of the other alleles also inhibited division (Table 1). C) FtsL, residues 54-99, was modelled (for illustration purposes) as an alpha helix since it is thought to form a continuous alpha helix with the TM and this region is also thought to form a coiled coil with FtsB. Altering residues in green leads to activation mutations, whereas altering those residues in red are dominant negative. Altering residues in yellow had no effect. Note that the activation mutations affect residues on one side of the helix whereas the dominant negative mutations affect residues on the other side. The red residues (including L86 and E87) identify a region designated AWI (<u>A</u>ctivation of Fts<u>WI</u>). The position of residues 24 and 28 in the cytoplasmic domain are indicated along with the transmembrane (TM) domain. The cytoplasmic domain of FtsL is required to recruit FtsW which in turn recruits FtsI. D) Cartoons depicting the effect of various mutations on the activation of FtsWI according to the model. Top, FtsN action makes AWI available; middle, FtsL^E88K^ is less dependent upon FtsN as the E88K substitution makes AWI available; bottom, FtsL^E87K^ is resistant to FtsN action and AWI does not become available or is defective in interaction with FtsWI.

Of residues comprising the CCD domain of FtsL residue E88 is the most conserved and mutational analysis indicated that loss of the negative charge results in the activation phenotype (10). The neighboring residue E87 is even more conserved (Fig. S1B) and was altered in one of our dominant negative mutants. Additional analysis indicates that changing this residue to amino acids other than aspartate produces a dominant negative phenotype (Fig. S2C). Thus, the loss of the negative charge in two neighboring glutamate residues yields contrasting phenotypes. Since loss of the negative charge in each case produced their respective phenotypes, it strongly suggests that these mutations disrupt rather than enhance interactions.

In our random mutagenesis screen, we did not isolate dominant negative mutations in *ftsB*, however, since six of the dominant negative mutations in *ftsL* overlapped the CCD, we used site-directed mutagenesis to alter the more conserved residues that overlap FtsB’s CCD domain. Seven residues flanking the CCD domain were altered, but none produced a dominant negative phenotype (Table 1). Six of these still complemented a *ftsB* deletion strain. This result suggests that the dominant negative mutations are unique to *ftsL*.

### Dominant negative FtsL mutants are defective in activation of septal PG synthesis

A dominant negative phenotype could result from incorporation of an FtsL mutant into the FtsQLB complex that fails to: 1) recruit downstream proteins (FtsWI), 2) respond to FtsN (FtsQLB locked in OFF state), or 3) generate an output signal in response to FtsN (ON state but fails to interact with a downstream partner). To test the first possibility, we assessed the localization of GFP-FtsI, which depends upon FtsW (3, 23). It was present in crossbands within filamentous cells following expression of *ftsL*^*E87K*^ or *ftsL*^*A90E*^ indicating recruitment to the Z ring was not an issue (Fig. S1D). Thus, the *ftsL* mutations either block the response to FtsN or a downstream event such as interaction with FtsWI.

The dominant negative *ftsL* mutations were tested to see if they could be rescued by a strong activation mutation (*ftsL*^*E88K*^) in *cis*. While *ftsL*^*R61E*^ and *ftsL*^*A90E*^ were readily rescued by *ftsL*^*E88K*^, f*tsL*^*L86F*^ and *ftsL*^*E87K*^ were not (Fig. S2A). If we assume that *ftsL*^*E88K*^ mimics FtsN action and switches FtsQLB to the ON state, it suggests that *ftsL*^*R61E*^ and *ftsL*^*A90E*^ are able to carry out steps downstream of FtsN action. Based on these results we suspected overexpression of *ftsN* would also rescue *ftsL*^*A90E*^ but not *ftsL*^*E87K*^. As expected overexpression of *ftsN* rescued *ftsL*^*A90E*^ but not *ftsL*^*E87K*^ (Fig. S2B). Since *ftsL*^*R61E*^ and *ftsL*^*A90E*^ can be rescued by enhancing the activation signal (by introducing an *ftsL* activation mutation or *ftsN* overexpression), it suggests they favor the OFF state but can carry out downstream events when activated. We therefore focused on f*tsL*^*L86F*^ and *ftsL*^*E87K*^ since it is unclear if they are locked in the OFF state or are unable to produce a signal in response to FtsN.

### Dominant negative FtsL mutants are rescued by FtsW activation mutants

Based on our results we hypothesized that activation of FtsWI requires a signal from the periplasmic domain of FtsL (AWI domain) which is made available by FtsN action or *ftsL* activation mutations. Activation alleles of *ftsW* might rescue a strong dominant negative *ftsL* allele since they require less input from FtsN. Two such *ftsW* alleles exist: *ftsW*^*M269I*^ (which weakly bypasses *ftsN* (12)) and *ftsW*^*E289G*^, which was isolated as described in the Materials and Methods and bypasses *ftsN*. The same mutation was recently isolated using another approach and also shown to bypass *ftsN* (biorivx 850073).

To see if these *ftsW* alleles could rescue f*tsL*^*L86F*^ or *ftsL*^*E87K*^, a plasmid with these alleles under an arabinose-inducible promoter (derivatives of pSD296 [P_*ara*_::*ftsL*]), as well as a compatible plasmid with *ftsW* alleles under an IPTG-inducible promoter (derivatives of pSEB429 [P_*204*_::*ftsW*]), were introduced into SD399 (*ftsL::kan*/pSD256 [*repA*^ts^::*ftsL*]). The resultant strains were spot tested at 37°C to deplete WT *ftsL* and arabinose and IPTG were added to induce the *ftsL* and *ftsW* alleles, respectively. Expression of *ftsW*^*M269I*^ and *ftsW*^*E289G*^, but not *ftsW*, rescued the dominant negative *ftsL* alleles (Fig. 3). These *ftsW* activation alleles still required the presence of *ftsL* as they could not bypass it (Fig. 3, right panel). Also, *ftsW*^*M269I*^ was able to rescue an allele containing both mutations (*ftsL*^*L86F*/*E87K*^) whereas overexpression of *ftsN* could not (Fig. S3A). These results indicate that *ftsL*^*L86F*/*E87K*^ cannot transmit the periplasmic signal in response to FtsN.

The above results demonstrate that the two dominant negative mutations (*ftsL* ^*L86F*^ or *ftsL*^*E87K*^, alone or combined) block FtsN but do not distinguish between whether they lock FtsQLB in the OFF state or prevent a downstream step (e.g, interaction with FtsWI). We suspect the latter for the following reasons. To rescue *ftsL* ^*L86F*^ or *ftsL*^*E87K*^, *ftsW*^*E289G*^ had to be overexpressed whereas the chromosomal level of *ftsW*^*E289G*^ is sufficient to bypass *ftsN* (expression of *ftsW* or the activation alleles complement an *ftsW* depletion mutant in the absence of IPTG [Fig. S4A] whereas 15 to 30 μM is required to rescue *ftsL* ^*L86F*^ or *ftsL*^*E87K*^). Consistent with this, expression of *ftsL*^*E87K*^ is toxic to a strain with *ftsW*^*M269I*^ on the chromosome (Fig. S4B) highlighting that an active *ftsW* allele cannot bypass the dominant negative *ftsL* mutation at the chromosomal level. Therefore, the results suggest that the dominant negative *ftsL* mutants are defective in interaction with FtsWI in the periplasm (lack of the periplasmic interaction may necessitate overexpression of *ftsW*^*M269I*^). Consistent with the *ftsL* mutations blocking a step downstream of FtsN action, an active *ftsB* mutation, *ftsB*^*E56A*^, which can also bypass *ftsN* (10), cannot suppress *ftsL*^*E87K*^ (Fig. S3B). This conclusion is also consistent with an activation mutation in *ftsL* or overexpression of *ftsN* being unable to rescue *ftsL*^*E87K*^ (Fig. S3A). Also, all substitutions in *ftsL*^*E87*^ that remove the negative charge are dominant negative (Fig. S1C) suggesting they disrupt, rather than enhance an interaction. We therefore favor the idea that these mutations in the AWI domain abrogate FtsL’s interaction with FtsWI and that under physiological conditions the AWI domain of FtsL in the ON state synergizes with ^cyto^FtsL to interact with FtsWI.

**Fig. 3.**
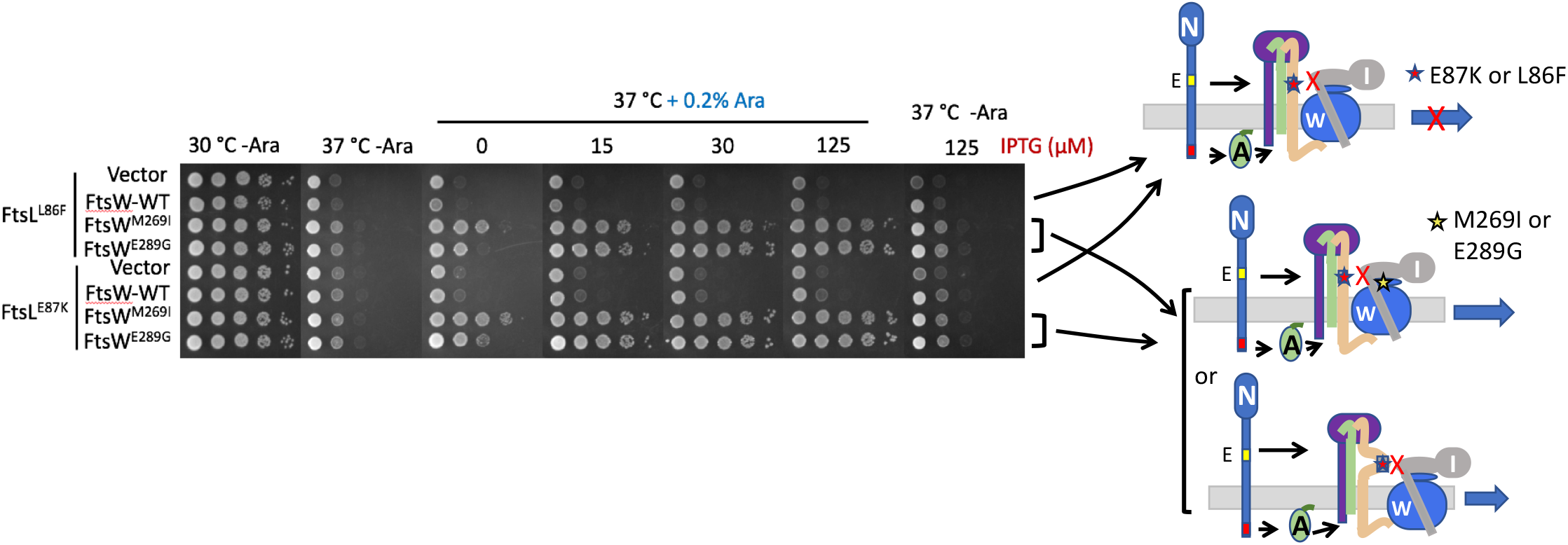
Response of dominant negative mutations in *ftsL* to FtsN and FtsW. Rescue of *ftsL* dominant negative mutations by overexpression of active *ftsW* mutants. SD399 (*ftsL::kan*/pSD256 [*repA*^ts^::*ftsL*]) containing derivatives of pSD296 (P_*ara*_::*ftsL*) with different alleles of *ftsL* was transformed with derivatives of pSEB429 (P_204_::*ftsW*) carrying WT *ftsW* or either of two active alleles of *ftsW*. Transformants were spot tested at 37°C (to deplete WT FtsL) in the presence of arabinose (to induce the *ftsL* allele present on derivatives of pSD296) and increasing concentrations of IPTG to induce alleles of *ftsW* (*ftsW, ftsW*^*M269I*^ or *ftsW*^*E289G*^). The panel on the far right reveals that these activated *ftsW* alleles cannot bypass *ftsL*. The cartoons depict interpretation of the results. In the upper cartoon the dominant negative *ftsL* mutations prevent FtsWI activation whereas the activated *ftsW* alleles do not require the input from FtsL.

### Loss of ^cyto^FtsL function rescued by activation mutations in the CCD domain of FtsL

One mutation from the random mutagenesis screen altered a residue in the ^cyto^FtsL domain (*ftsL*^*L24K*^). Although weak, adding a second mutation altering a highly conserved residue in this domain (*ftsL*^*I28K*^) yielded a stronger dominant negative phenotype (Fig. S5A). Since this domain of FtsL is required for FtsW recruitment (13), it suggested that FtsL^L24K^, FtsL^I28K^ and the double mutant assemble into a complex with FtsQB that poorly recruits FtsW. Consistent with this, deletion of the cytoplasmic domain of FtsL (FtsL^Δ1-30^) also produced a strong dominant negative phenotype (Fig. S5B).

Since FtsN is proposed to switch FtsQLB to the ON state to activate FtsWI (10, 11), we speculated above that this switch involves a conformational change that exposes AWI to activate FtsWI. If this is the case, then the activation mutations may compensate for the loss of ^cyto^FtsL by making the AWI domain available to recruit FtsWI as well as to activate it. As expected *ftsL*^Δ1-30^ failed to complement Δ*ftsL*, however, *ftsL*^Δ*1-30*^ carrying two activation mutations (*ftsL*^*G92D*^ and *ftsL*^*E88K*^) restored colony formation indicating that both recruitment and activation of FtsW were restored (Fig. 4A). Further tests showed that both activation mutations were required for rescue (Fig. S6A). The rescue was fairly effective as the average cell length of the strain expressing *ftsL*^Δ1-30/*G92D/E88K*^ was only twice that of a strain expressing *ftsL* (Fig. S6B), whereas, the strain expressing *ftsL*^Δ1-30^ was extremely filamentous. Combining these two activation mutations also eliminated the toxicity of the *ftsL*^*L24K/I28K*^ allele (Fig. S6C) and rescued its ability to complement (Fig. S6D). These results are consistent with a model in which the *ftsL* activation mutations cause a conformational change in FtsQLB that makes AWI available to recruit and activate FtsWI. It follows that under physiological conditions the arrival of FtsN results in the exposure of ^AWI^FtsL which cooperates with ^cyto^FtsL to recruit and activate FtsWI.

**Fig. 4.**
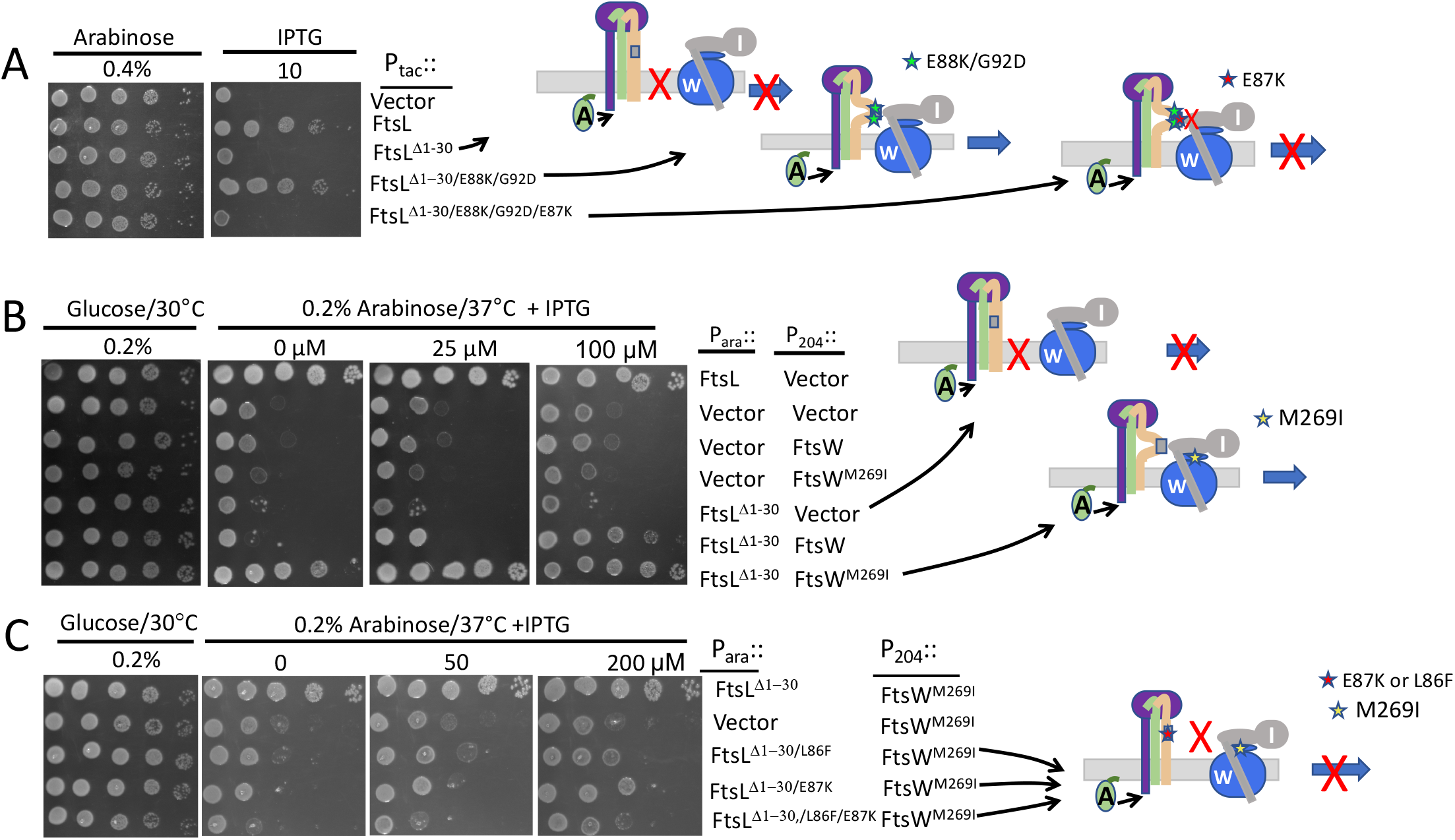
Loss of FtsL^cyto^ function rescued by activation mutations. Effect of *ftsL* activation and dominant negative mutations on the rescue of FtsL^Δ1-30^. A) *ftsL*^*Δ1-30*^ is rescued by *ftsL* activation mutations which is negated by an *ftsL* dominant negative mutation. SD439 (*ftsL::kan*/pSD296 [P_*ara*_::*ftsL*]) was transformed with derivatives of pKTP105 (*P*_*T5*_::*ftsL*) carrying various alleles of *ftsL* inducible with IPTG. The strains were spotted on plates without arabinose (to deplete WT *ftsL*) but with IPTG (to induce the various alleles of *ftsL* present in derivatives of pKTP105). The cartoons to the right depict the results. B) Overexpression of *ftsW*^*M269I*^ rescues *ftsL*^*Δ1-30*^. SD399 (*ftsL::kan*/pSD256 [*repA*^TS^ P_syn135_::*ftsL*]) carrying pKTP107 (P_ara_::*ftsL*^*Δ1-30*^) was transformed with compatible plasmids expressing different alleles of *ftsW* (derivatives of pSEB429 [P_204_::*ftsW*]) under control of and IPTG inducible promoter. Transformants were spotted on plates at 37°C (to deplete WT *ftsL*) in the presence of 0.2% arabinose to induce *ftsL* alleles and increasing concentrations of IPTG to induce *ftsW* alleles. The cartoon indicates that FtsW^M269I^ is recruited by FtsL^*Δ1-*30^. C) Dominant negative *ftsL* mutations negate rescue of *ftsL*^*Δ1-30*^ by *ftsW*^*M269I*^. Strain SD399 (*ftsL::kan*/pSD256 [*repA*^ts^-*ftsL*]) was transformed with a plasmid (derivatives of pKTP107 [P_ara_::*ftsL*^*Δ1-30*^]) with *ftsL*^*Δ1-30*^ under arabinose promoter control and a compatible plasmid (pSEB429 [P_204_:: *ftsW*]) that carries *ftsW* or *ftsW*^*M269I*^ under control of an IPTG-inducible promoter. The presence of a dominant negative *ftsL* mutation negates rescue by the activated FtsW.

Since the *ftsL* activation mutations appear to mimic FtsN action, we expected that overexpression of *ftsN* would also rescue *ftsL*^Δ*1-30*^. To test this, an *ftsL* depletion strain was transformed with a plasmid expressing *ftsL*^Δ*1-30*^ and a plasmid that overexpresses *ftsN* to a level that is sufficient to bypass *zipA* or *ftsEX* (21). The increased FtsN rescued *ftsL*^Δ*1-30*^ (Fig. S6E) suggesting that the AWI became available to recruit and activate FtsWI. Thus, overexpressing *ftsN* is comparable to combining the two activation mutations (*ftsL*^*G92D*^ and *ftsL*^*E88K*^) in rescuing *ftsL*^Δ*1-30*^.

### Dominant negative *ftsL* mutations negate rescue by activation mutations

If the *ftsL* activation mutations rescue *ftsL*^Δ*1-30*^ by making AWI available to recruit and activate FtsWI, the dominant negative mutations should impair rescue by blocking the interaction. As seen in Fig. 4A, addition of *ftsL*^*E87K*^ negated the rescue of *ftsL*^Δ1-30^ by the activation mutations. This result is consistent with *ftsL*^*E87K*^ blocking interaction between the AWI domain and FtsWI.

The FtsQLB complex probably exists in equilibrium between ON and OFF states with the activation mutations and overexpression of FtsN favoring the ON state (AWI available). Overexpression of FtsW or FtsW ^M269I^ may also tip the equilibrium to the ON state and rescue *ftsL*^Δ*1-30*^ as the increased level of FtsW may promote capture of the ON state. Indeed, expression of *ftsW*^*M269I*^, even at low levels of induction, rescued *ftsL*^Δ*1-30*^ and at higher levels of induction WT *ftsW* also started to rescue (Fig. 4B).

Earlier we showed that overexpression of *ftsW*^*M269I*^ and *ftsW*^*E289G*^ but not *ftsW* rescued *ftsL* carrying dominant negative mutations (Fig. 3). This result is consistent with these activated mutants being recruited by the FtsL mutants (through ^cyto^FtsL) but not requiring an activation signal from the AWI domain (via FtsN) (12). In the absence of ^cyto^FtsL, however, our results suggest rescue is dependent upon a functional AWI in FtsL^peri^. If so, the dominant negative mutations should be detrimental in this context. We found that the addition of either of two dominant negative mutations (*ftsL*^*L86F*^ or *ftsL*^*E87K*^) to *ftsL*^Δ1-30^ prevented rescue by FtsW^M269*I*^ (Fig. 4C). These results are consistent with AWI being required to recruit FtsWI in the absence of ^cyto^FtsL. It is worth noting that when either of two FtsL domains is nonfunctional (either inactivation of the cytoplasmic domain or the presence of the dominant negative mutations [such as L86F and E87K] in full length FtsL) the active FtsW mutants must be overexpressed to rescue growth (see Discussion).

### Rescue of FtsL^Δ1-30^ by overexpression of FtsI

In the hierarchical assembly pathway, FtsW is recruited in a ^cyto^FtsL dependent manner and FtsI is then recruited by interaction between FtsW and the transmembrane segment of FtsI (23). However, we considered the possibility that with FtsL^Δ1-30^ the recruitment is reversed or FtsWI is recruited as a complex through interaction of AWI with FtsI. This thinking was driven in part by geometric constraints. The periplasmic domain of FtsL is thought to be a continuous alpha helix with its transmembrane domain such that the AWI domain would extend about ∼45Å away from the cytoplasmic membrane (15) (Fig. S7). In the RodA-PBP2 structure (homologous to FtsW-FtsI) the non-penicillin binding domain of PBP2 rests on top of RodA and extends into the periplasm (24). Assuming FtsW-FtsI adopts a similar structure, FtsI could contact AWI.

If FtsI interacts with the AWI domain, overexpression of *ftsI* may rescue FtsL^Δ1-30^ by increasing the interaction and shifting the equilibrium of FtsQLB from OFF to ON through mass action. To test this, we compared the ability of the overexpression of *ftsI* and *ftsW* to rescue FtsL^Δ1-30^. As shown in Fig. 5A, expression of *ftsI* was much more efficient than *ftsW* in rescuing FtsL^Δ1-30^. The efficient rescue of FtsL^Δ1-30^ by FtsI suggests that it contacts AWI and converts FtsL^Δ1-30^ into an active form similar to the activation mutations (Fig. 4A). The rescue of FtsL^Δ1-30^ by overexpression of FtsW may involve formation of a FtsWI complex that interacts with AWI. The efficient rescue of FtsL^Δ1-30^ by activated FtsW (compared to WT FtsW seen in Fig. 4B) may be due to it more readily forming a complex with FtsI.

**Fig. 5.**
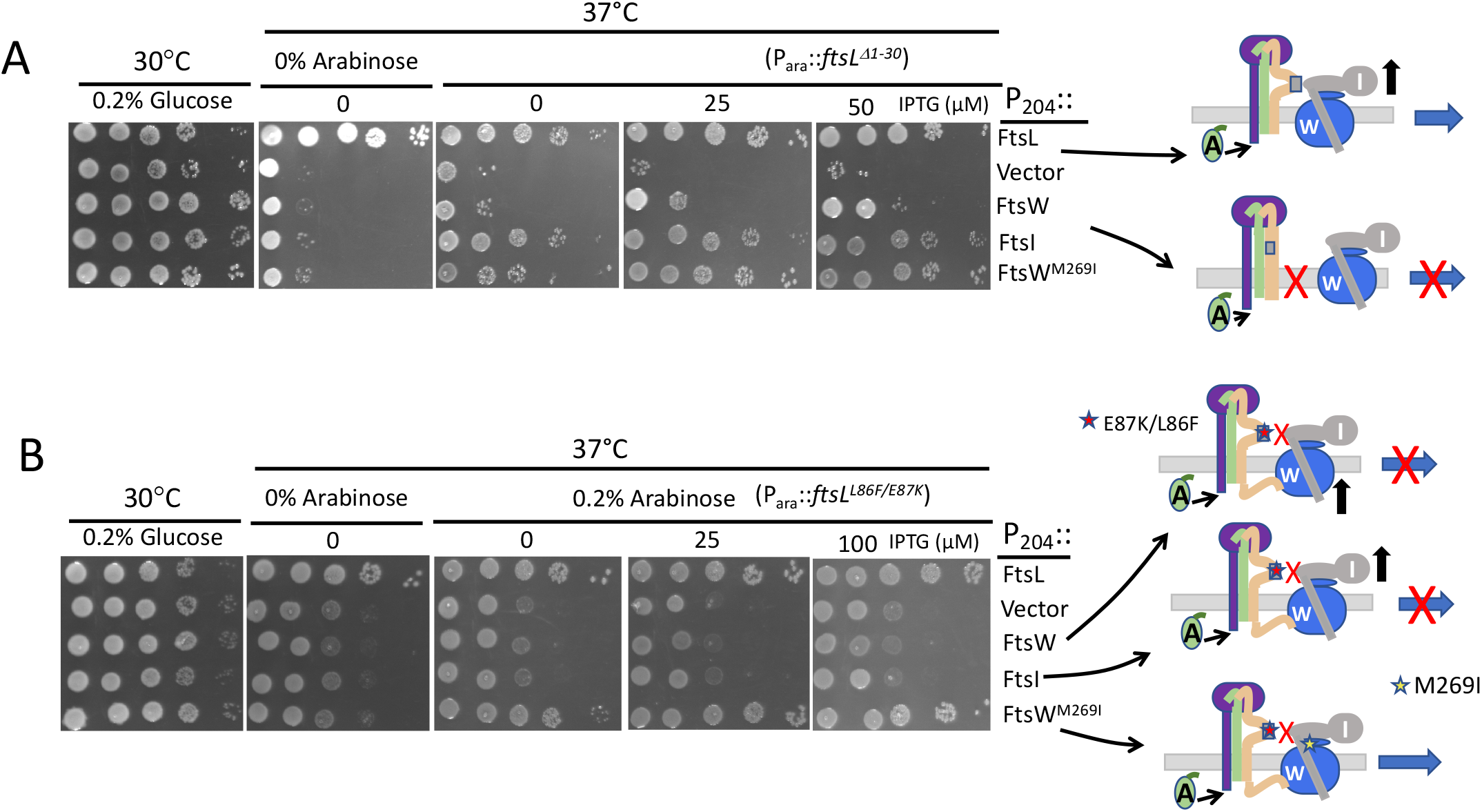
Rescue of FtsL^Δ1-30^ by *ftsI* overexpression. A) *ftsI* overexpression rescues FtsL^Δ1-30^. To test if overexpression of *ftsI* could rescue FtsL^*Δ*1-30^, PK4-1 (*ftsL::kan*/pKTP108 [*repA*^*TS*^::*ftsL*]) was transformed with a plasmid expressing *ftsL*^Δ1-30^ (pKTP107X/P_ara_::*ftsL*^*Δ*1-30^-*6Xhis*) and a plasmid expressing *ftsI* (pSEB420/P_204_::*ftsI*) or *ftsW* (pSEB429/P_204_::*ftsW*) under an IPTG-inducible promoter. The strains were spot tested on plates at 37°C to deplete *ftsL*. In addition, arabinose was added to induce *ftsL*^*Δ1-30*^ and increasing concentrations of IPTG were added to induced *ftsI* or *ftsW*. B) *ftsI* overexpression cannot rescue a dominant negative *ftsL* allele. SD399 (*ftsL::kan*/pSD256 [*repA*^ts^::*ftsL*]) was transformed with pSD296-2 (P_ara_::*ftsL*^*L86F/E87K*^) and pSEB420 (P_204_::*ftsI*) or pSEB429 (P_204_::*ftsW*). Transformants were spot tested at 37°C (to deplete *ftsL*) on plates containing 0.2% arabinose (to induce *ftsL*^*L86F/E87K*^) and increasing concentrations of IPTG to induce *ftsI* or *ftsW*.

The above results indicate that the signal from FtsN via the AWI domain goes through FtsI. As shown earlier, expression of activated alleles of *ftsW* suppresses *ftsL*^*L86F*^ or *ftsK*^*E87K*^ as they no longer require the signal from AWI. In contrast, WT *ftsW* cannot suppress these alleles as it still requires the AWI activation signal. Likewise, overexpression of *ftsI* should not rescue full length FtsL carrying the dominant negative *ftsL* mutations. As expected, overexpression of *ftsI* was unable to suppress *ftsL*^*L86F*/*E87K*^ indicating the AWI signal was still required (Fig. 5B).

The possibility that AWI recruits and activates FtsWI by acting through FtsI was further examined by testing FtsI mutants isolated by the Weiss lab (25). These mutants localize to the division site but fail to complement and recruit FtsN. We thought it possible that some of these FtsI mutants were defective in relaying an activation signal from AWI to FtsW, and thus unable to initiate a positive feedback loop leading to FtsN accumulation. Therefore, we compared the rescue of these FtsI mutants by active mutants of FtsL and FtsW (FtsL^G92D/E88K^ and FtsW^M269I^, respectively) which are less dependent upon FtsN. The rationale was that if an active FtsL converts FtsW to an active form then the rescue of FtsI mutants by FtsW^M269I^ should be equal to or greater than that by an active FtsL. On the other hand, if FtsL acts through FtsI to activate FtsW, then FtsL^G92D/E88K^ may be able to rescue FtsI mutants more efficiently than FtsW^M269I^. Indeed, of seven FtsI mutants tested, two mutants (FtsI^S61F^ and FtsI^R210C^) were rescued by both FtsW^M269I^ and FtsL^G92D/E88K^ (Fig. 6 and S8). However, FtsL^G92D/E88K^ rescued two additional mutants (FtsI^G57D^ and FtsI^V86E^) not rescued by FtsW^M269I^. The rescue of these two mutants by an activated FtsL (but not an activated FtsW) suggests that AWI acts through FtsI to activate FtsW rather than acting directly on FtsW.

**Fig. 6.**
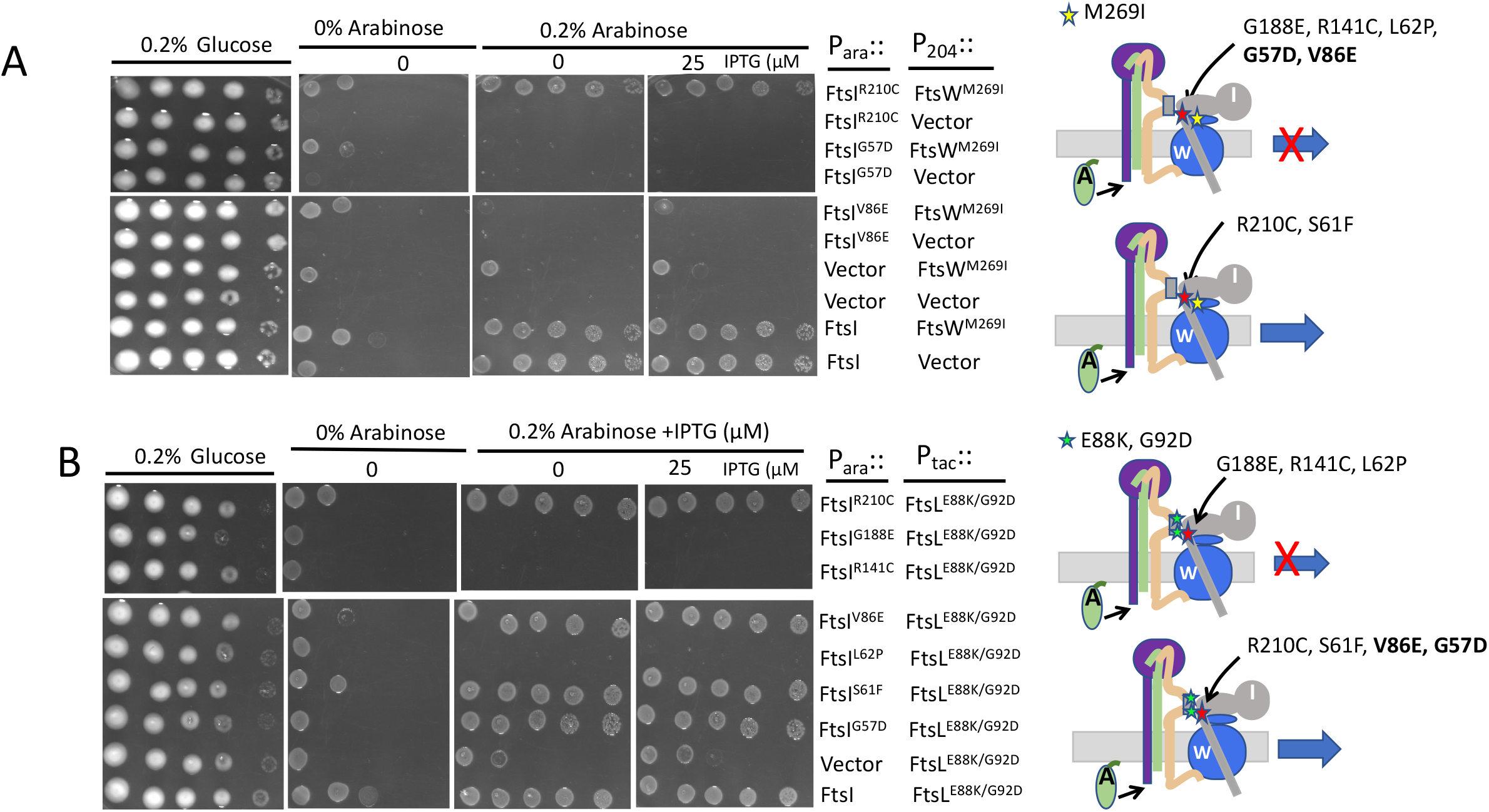
Rescue of FtsI mutants by activated FtsL and FtsW mutants. A) Rescue of FtsI mutants by FtsW^M269I^. To test if the FtsI mutants could be rescued by an activated allele of *ftsW*, MCI23 (*ftsI23*^*ts*^ *recA::spc)* was transformed with compatible plasmids expressing an activated allele of *ftsW* (pSEB429 [P_*204*_::*ftsW*^*M269I*^]) and *ftsI* alleles under arabinose promoter control (derivatives of pKTP109 [P_ara_::*ftsI*]). Transformants were spot tested on plates at 37°C (to inactivate *ftsI23*^ts^) and arabinose added to induce the *ftsI* alleles and increasing concentrations of IPTG to induce *ftsW*^*M269I*^. Note: additional alleles of *ftsI* were not rescued by *ftsW*^*M269I*^ (Fig. S8). B) Rescue of FtsI mutants by *ftsL*^*E88K/G92D*^. To test rescue of FtsI mutants by activated FtsL, MCI23 (*ftsI23*^*ts*^ *recA::spc)* was transformed with compatible plasmids expressing an activated allele of *ftsL* (pKTP100* [P_*tac*_::*ftsL*^*E88K/G92D*^]) and the various *ftsI* alleles under arabinose promoter control (derivatives of pKTP109 [P_ara_::*ftsI*]). Transformants were spot tested on plates at 37°C (to inactivate *ftsI23*^ts^) and arabinose added to induce the *ftsI* alleles and increasing concentrations of IPTG to induce *ftsL*^*E88K/G92D*^.

### Interaction between FtsL and FtsWI

Our results are consistent with an interaction between the cytoplasmic domain of FtsL and FtsW leading to recruitment of FtsWI and between the periplasmic domain of FtsL with FtsI which is required for activation of FtsWI. As shown above, when the ^cyto^FtsL-FtsW interaction is eliminated activation mutations in *ftsL* and *ftsW* are able to rescue division which is abrogated by the dominant negative mutations in *ftsL*. To obtain additional support for interactions between the various proteins we tested the effect of these mutations using the bacterial two hybrid (BTH) system. We observed strong interactions between FtsL and FtsW and between FtsL and FtsI, however, which were eliminated when the cytoplasmic domain of FtsL was deleted, consistent with ^cyto^FtsL involved in recruiting FtsWI (FtsL^Δ1-30^, Fig. 7A). This allowed us to use FtsL^Δ1-30^ to assess the effects of the activation mutations in *ftsL* and *ftsW*. Although the *ftsW* activation mutation had little effect, the addition of two *ftsL* activation mutations resulted in a strong interaction between FtsL^Δ1-30^ and FtsI and a weaker interaction between FtsL^Δ1-30^ and FtsW (Fig. 7B). The weak interaction with FtsW suggests that FtsI was an intermediate. Importantly, the addition of a dominant negative mutation (*ftsL*^*E87K*^) eliminated the interaction conferred by the activation mutations. The effects of these *ftsL* mutations in the BTH correlate with the effects these mutations have on the rescue of FtsL^Δ1-30^; the *ftsL* activation mutations promote rescue and interaction which is negated by an *ftsL* dominant negative mutation (Fig. 4A).

**Fig. 7.**
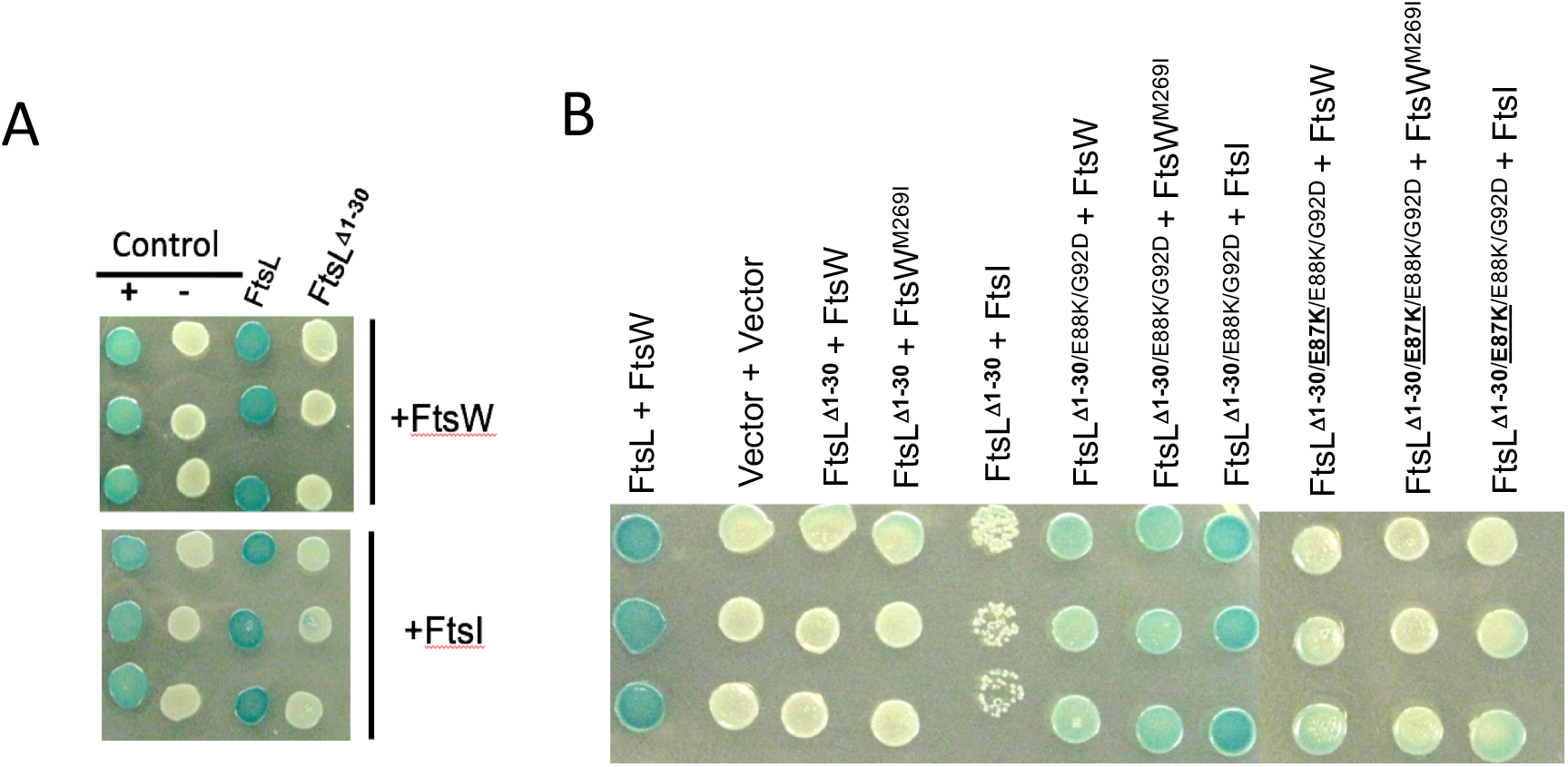
Interaction between FtsL and FtsWI assessed with the BTH system. A) Effect of ^cyto^FtsL on the interaction between FtsL and FtsWI. Strain DMH1 was transformed with plasmids carrying various alleles of *ftsL* (pUC18 derivatives) and plasmids expressing *ftsW* or *ftsI* (pKT25 derivatives). Three transformants were picked and spot tested for each pair of constructs. The positive control contained plasmids pUC18C-zip and pKT25-zip whereas the negative control contained the corresponding empty vectors. B) The effect of activation and dominant negative mutations in *ftsL* on the interaction of FtsL^Δ1-30^ with FtsW and FtsI. Three transformants from each transformation of DHM1 with plasmids carrying the f*tsL* and *ftsW* or *ftsI* alleles were spotted on plates containing the color indicator. The *ftsL* alleles were contained in pKT25 and the *ftsW* and *ftsI* alleles were in pUT18. The plates were photographed after overnight incubation.

### Rescue of Δ*ftsL* by MalF-FtsL and FtsW-FtsK fusions

Next, we tested if the periplasmic portion of FtsL transported to the periplasm could activate FtsWI in the absence of full length FtsL. To do this, a MalF-FtsL fusion was constructed under the control of an IPTG-inducible promoter in which the cytoplasmic and TM domains of FtsL were replaced with the corresponding regions of MalF (^cyo/TM^MalF-^peri^FtsL). In contrast to FtsL^*Δ*1-30^, this MalF-FtsL fusion was not dominant negative (Fig. S9A) indicating the TM region of FtsL must be present for the fusion to displace FtsL from the FtsQLB complex and disrupt FtsW recruitment. This is consistent with the transmembrane (TM) region of FtsL being unique (26) and the TMs of FtsL and FtsB being required for these proteins to interact (16, 18). Furthermore, the MalF-FtsL fusion was unable to complement an *ftsL* depletion strain even if the strain carried a *ftsW*^*M269I*^ mutation and the *ftsL* construct carried the two activation mutations (Fig. S9B).

Since FtsQLB is required to recruit FtsWI and the MalF-FtsL fusion is insufficient to recruit FtsW, we used an FtsW-^cyto^FtsK fusion which complements a *ftsK* deletion mutant, as well as a *ftsW* deletion mutant, indicating it bypasses FtsQLB for recruitment (27) (and data not shown). This MalF-FtsL fusion was unable to replace FtsL and rescue the growth of a strain containing FtsW-^cyto^FtsK, even if the fusion carried activation both *ftsL* activation mutations (Fig. 8). This inability to activate the FtsW-^cyto^FtsK fusion could be for a variety of reasons including that FtsQB is uncoupled from FtsL and the FtsW-^cyto^FtsK likely competes with endogenous FtsW for FtsI. However, this MalF-FtsL fusion with the two activation mutations was able to rescue a strain containing the FtsW-^cyto^FtsK fusion with the *ftsW*^*M269I*^ mutation. (Fig. 8). Even the MalF-FtsL fusion without the *ftsL* activation mutations partially rescued growth at higher induction levels. These results suggest that MalF-FtsL acts on FtsI associated with the FtsW^M269I-cyto^FtsK fusion that is already at the Z ring to rescue growth. Since the activation mutations in *ftsL* potentiate MalF-FtsL, it suggests that in addition to making AWI available within the FtsQLB complex, they may also alter the structure of AWI to enhance its interaction with FtsWI (since MalF-FtsL is not part of the FtsQLB complex).

**Fig. 8.**
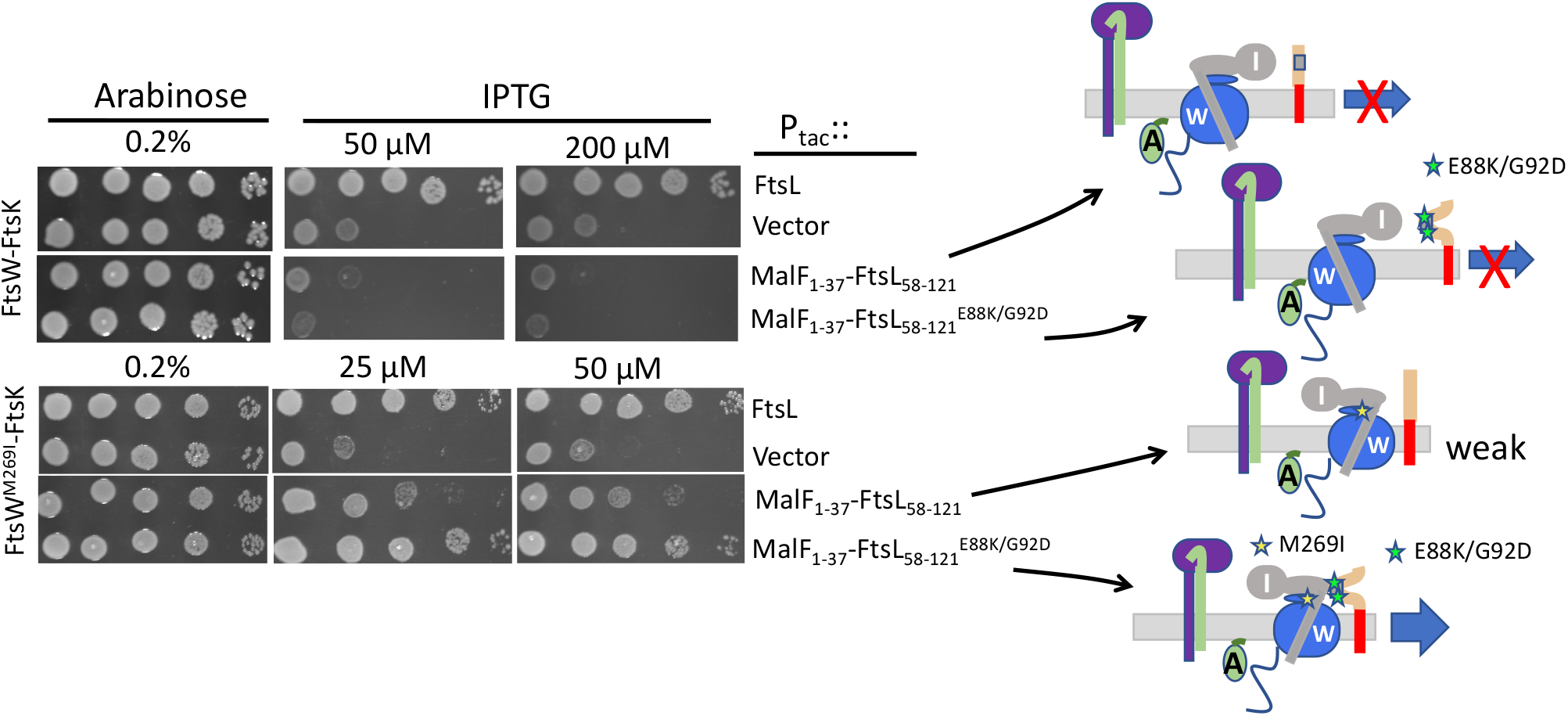
Rescue of a FtsW-FtsK fusion by a MalF-FtsL fusion. Activation mutations allow a *malF-ftsL* fusion to complement *ΔftsL* in the presence of a *ftsW-*^*cyto*^*ftsK* fusion. Plasmids pKTP103 (P_*tac*_::*malF*^*1-37*^*-ftsL*^*58-121*^*-6Xhis*) was introduced into a *ftsL* depletion strain (SD439 *ftsL::kan*/pSD296 [P_*ara*_::*ftsL*]) in the presence of a plasmid constitutively expressing a FtsW-FtsK^cyto^ fusion without or with an activation mutation (pND16 [P*ftsK::ftsW*-^*cyto*^*ftsK*] or pND16* [P*ftsK:: ftsW*^*M269I*^-^*cyto*^*ftsK*], respectively). The strains were spot tested on plates without arabinose (to deplete WT *ftsL*) and in the presence of IPTG to induce *malF-ftsL* (with or without the activation mutations [*ftsL*^*E88K*^ and *ftsL*^*G92D*^]). The cartoon to the right depicts the activity of the FtsL constructs.

## Discussion

Here we investigated how septal PG synthesis in the divisome is activated by FtsN and identified a critical and unique role for FtsL. Our results are consistent with the recruitment of FtsW requiring the cytoplasmic domain of FtsL and the activation of FtsWI being dependent upon AWI in the periplasmic domain of FtsL. Based upon the seminal work by the de Boer lab, which is supported by the work from the Bernhardt lab (10-11) and our results (12) and those herein, we propose that the arrival of FtsN leads to a conformational change in the FtsQLB complex that makes the AWI domain of FtsL, as defined by the dominant negative *ftsL* mutations, available to activate FtsWI by acting through FtsI. Activation mutations in the CCD domain of FtsL as well as those in FtsB mimic FtsN action to cause a conformational change in FtsQLB to expose the AWI domain. This model is supported by the ability of activation mutations in *ftsL* to rescue FtsL mutations (FtsL^Δ1-30^ and FtsL^L24K/I28K^) deficient in FtsWI recruitment and by the dominant negative mutations in *ftsL* (*ftsL*^*L86F/E87K*^) negating the rescue. The effects of these *ftsL* mutations (both activation and dominant negative) on the rescue of the FtsL mutants correlates with their effects on the observed interaction between FtsL and FtsWI in the BTH. The model is also supported by the ability of the expression of *ftsI* to rescue FtsL^Δ1-30^ far more efficiently than *ftsW*. Furthermore, FtsL acting on FtsI to activate FtsW is supported by the ability of an activated FtsL mutant to rescue FtsI mutants not rescued by an activated FtsW. Thus, we propose that as a result of ^E^FtsN action the AWI domain of FtsL becomes available to interact with FtsI within the FtsWI complex to activate FtsWI and synergizes with ^cyto^FtsL in stabilizing the FtsWI complex in the divisome.

### The AWI domain

Altering seven residues in the periplasmic domain of FtsL produced a dominant negative phenotype. All, except for one, are clustered together around the CCD. Among them we focused on L86 and E87 and believe these are central to the AWI domain. This suggestion is based upon: 1) L86 and E87 are conserved and loss of the negative charge at E87 is sufficient to produce a dominant negative allele (suggesting disruption of an interaction), 2) the *ftsL*^*L86F*^ or *ftsL*^*E87K*^ dominant negative mutations are not suppressed by activation mutations (*ftsL*^*E88K*^ or *ftsB*^*E56A*^) or *ftsN* overexpression, 3) ^cyto^FtsL mutants that fail to recruit FtsWI are rescued by addition of two *ftsL* activation mutations (*ftsL*^*E88K/G92D*^), 4) the rescue of ^cyto^FtsL mutants by *ftsL* activation mutations or overexpression of *ftsW*^*M269I*^ is negated by adding dominant negative mutations (*ftsL*^*L86F*^ or *ftsL*^*E87K*^) and 5) the effects of these mutations on the interaction of FtsL with FtsW in the BTH correlate well with the effects of these mutations on the rescue of FtsL^Δ1-30^. It is likely that other regions of FtsL (and FtsB) such as the transmembrane domains (TM) and coiled coil domains are also involved in interaction with FtsWI, however, such interactions are not sufficient to support recruitment following the loss of ^cyto^FtsL. In support of this, we could bypass *ftsQ* with an activated allele of *ftsA* but were unable to bypass *ftsB* or *ftsL* (data not shown).

The dominant negative mutations in *ftsL* are less responsive to FtsN and most overlap the CCD domain which was defined by hyperactive mutations that are less dependent upon FtsN (10, 11). Despite the overlap, the residues comprising each domain mostly lie on opposites sides of a putative helix consistent (Fig. 2C). The dominant negative mutations appear to be unique to *ftsL* as were unable to isolate any such mutations in *ftsB*. Previous studies suggested that FtsN induces a change in FtsQLB from an OFF to ON conformation (10), however, it was not clear how this switch led to activation of FtsWI. Here we identify the AWI domain of FtsL and suggest that the function of the conformational switch is to make AWI available to interact with FtsWI. Since FtsQLB is likely a dimer, the conformational change may involve disruption of this dimer which makes AWI available, however, this will require further study (15, 16, 28).

Additional evidence for the unique importance of the periplasmic domain of FtsL comes from the ability of the MalF-^peri^FtsL fusion to rescue a FtsW-^cyto^FtsK fusion when both are carrying activation mutations. The FtsW-^cyto^FtsK fusion is unable to support growth as it is uncoupled from FtsQB due to the absence of full length FtsL. On the other hand, the MalF-^peri^FtsL fusion is not recruited to the divisome as it does not form a complex with FtsQB. Nonetheless, the ability of the MalF-^peri^FtsL to collaborate with FtsW-^cyto^FtsK (when both are carrying activation mutations) to rescue growth suggests that the periplasmic domain of FtsL acts on FtsW-^cyto^FtsK complexed with FtsI.

### Conditions that rescue FtsL^Δ1-30^ favor interaction between FtsL and FtsI

Surprisingly, loss of the cytoplasmic domain of FtsL, which prevents recruitment of FtsWI and blocks cell division, could be rescued by activation mutations in the periplasmic domain of FtsL as well as by overexpression of FtsN. We reasoned these activation conditions expose an interaction that normally occurs when the divisome is activated and that this interaction is able to compensate for the loss of ^cyto^FtsL. The finding that the these *ftsL* activation mutations to *ftsL*^Δ1-30^ promote interaction between FtsL and both FtsW and FtsI and that these interactions were negated by the addition of a dominant negative mutation (in AWI) supports this model. These results suggest that FtsL within the FtsQLB complex functions as a transmembrane clamp to stabilize the FtsWI complex within the divisome. The cytoplasmic domain of FtsL is required to recruit FtsW which in turn recruits FtsI. FtsN action then frees the AWI domain to interact with FtsI and as we have shown here, this domain when freed is able to rescue *ftsL*^Δ1-30^ indicating FtsWI recruitment is restored.

If FtsQLB exists in equilibrium between that ON and OFF states, we reasoned that expression of the downstream partner might also rescue *ftsL*^Δ1-30^ by capturing the ON form and pulling the equilibrium in that direction. In fact, the active form of FtsW was quite effective in rescuing *ftsL*^Δ1-30^, much more so than FtsW. However, expression of FtsI was very effective in rescuing *ftsL*^Δ1-30^ and much more so that overexpression of FtsW, which barely rescued at high overexpression. This suggested: 1) FtsI is the direct downstream target of AWI, 2) rescue by expression of FtsW likely involves formation of an FtsWI complex recruited by AWI, and 3) raises the possibility that the activated form of FtsW interacts more strongly with FtsI. Consistent with the rescue of *ftsL*^Δ1-30^ by expression of FtsI or activated FtsW being dependent upon the interaction of AWI with FtsWI in the periplasm, it was prevented by the addition of the dominant negative *ftsL* mutations. This is in stark contrast to the suppression of the dominant negative mutations in full length *ftsL* by activated FtsW. When full length FtsL is present, an FtsW activated by mutation is recruited normally and no longer requires the activation signal so the dominant negative mutations do not prevent the rescue (although rescue is aided by overexpression of the activated FtsW). On the other hand, FtsI cannot rescue as it is not activated and still depends upon the AWI signal.

Our results suggest that FtsWI forms a dynamic complex but it is the complex that is preferred by FtsL. If FtsWI formed a stable complex then overexpression of FtsW would be toxic as excess FtsW would titrate FtsI away from the division site inhibiting division. However, overexpression of *ftsW* rescued *ftsL*^Δ1-30^ and is not toxic in WT cells. Also, when FtsQLB is overexpressed and purified, FtsW and FtsI only copurify efficiently if they are both expressed indicating that the FtsWI complex interacts more stably with FtsQLB than FtsW or FtsI alone (29). Thus, overexpression of FtsW may favor complex formation with FtsI and septal localization to rescue *ftsL*^Δ1-30^. More efficient rescue by an activated FtsW could be due to it favoring complex formation with FtsI. On the other hand, the effect of FtsI expression on the rescue of *ftsL*^Δ1-30^ is probably distinct from complex formation with FtsW, likely involving a direct interaction with AWI, otherwise the rescue of *ftsL*^Δ1-30^ by FtsW and FtsI should be comparable, since overexpression of either should promote complex formation.

The product of FtsN action is an activated FtsWI complex in which both FtsW and FtsI are active. The ability of active FtsW mutants to suppress the dominant negative FtsL mutants (and bypass the periplasmic signal) indicates that an active FtsW leads to an active FtsWI complex. Among previously isolated FtsI mutants we found some that were rescued by both an active FtsW mutant and an active FtsL mutant. However, an activated FtsL was rescued two additional FtsI mutants that could not be rescued by an activated FtsW so that an active FtsWI complex is formed. This suggests that AWI acts on FtsI to activate FtsW and does not act directly on FtsW. In other words, the signal transmission from FtsN is from ^peri^FtsL→ FtsI→ FtsW and not ^peri^FtsL→ FtsW→ FtsI.

Although *in vitro* results suggest that FtsQLB acts as an inhibitor with FtsL inhibiting PBP1b and FtsQ inhibiting FtsI and therefore FtsW (29), our results are more compatible with a model in which AWI is sequestered within FtsQLB which becomes available upon FtsN action to activate FtsWI. The findings that *ftsL* activation mutations rescue FtsL^Δ1-30^ and promote interaction between FtsL^Δ1-30^ and FtsWI in the BTH is consistent the FtsL-FtsWI interaction activating FtsWI. This conclusion is also supported by the *ftsL* dominant negative mutations negating both of these activities.

### Comparison of models for divisome and elongasome activation

It is interesting to compare our model for FtsWI activation with the model proposed for activation of the RodA-PBP2 pair that are part of the elongasome (homologous to FtsW-FtsI [PBP3]). This model is based upon: 1) the structure of the MreC-PBP2 complex (30), and 2) the finding that mutations that bypass *mreC* and activate RodA-PBP2 map to the nonpenicillin or pedestal domain of PBP2 (31). It is thought that these mutations mimic the binding of MreC to PBP2 altering the conformation of PBP2 which results in the activation of RodA. In this way the activity of RodA and PBP2 are coupled to ensure RodA only makes glycan strands when its cognate PBP is present. This is remarkably similar to our model for FtsW-FtsI (PBP3) activation with FtsL being analogous to MreC. The isolation of FtsW activation mutants that bypass FtsN suggests that an activated FtsW results in an active FtsI. Furthermore, an active FtsW mutant can rescue dominant negative FtsL mutants (i.e. bypass the signal from FtsN) indicating FtsI is also activated. Thus, we propose that FtsN action alters the conformation of FtsQLB so that AWI becomes available to interact with FtsI leading to conformational change in FtsI that activates FtsWI’s enzymatic activities.

## Materials and Methods

### Bacterial Strains and Growth Conditions

Bacterial strains are listed in Table S1A. JS238 [*MC1061, araD* Δ(*ara leu) galU galK hsdS rpsL* Δ(*lacIOPZYA)X74 malP::lacI*^*Q*^ *srlC::*Tn*10 recA1*] was primarily used for screening for *ftsL* and *ftsB* dominant negative mutations and as a host for most cloning experiments. W3110 was used to generate SD399, SD439 and SD285. To construct SD399 [*W3110, ftsL::kan/*pSD256], P1 phage grown on BL156 [*ftsL::kan*/*pBL195*] was used to transduce *ftsL::kan* into W3110/pSD256 by selecting for Kan resistance on LB agar plates containing 25 µg/ml kanamycin, 50 µg/ml spectinomycin and 8 mM sodium citrate at 30°C. Several colonies were subcloned onto fresh plates of the same composition at 30°C and were further screened for temperature sensitivity at 42°C. SD439 was created by transforming SD399 with pSD296 (P_*ara*_::*ftsL*) and selecting the transformants that grow at 42°C (to remove pSD256) in the presence of 10 µg/ml chloramphenicol and 0.2% arabinose. Colonies were streaked and further tested for spectinomycin sensitivity (indicating loss of pSD256). Construction of SD285 [*leu::*Tn*10 bla lacI*^q^ *P*_207_-*gfp-ftsI*] involved transduction with P1 phage grown on EC436 [MC4100 *Δ*(λ*attL-lom*)::*bla lacI*^q^ *P*_207_-*gfp-ftsI*] into S3 (W3110 *leu::*Tn*10)*. Transductants were selected in LB agar plates containing 25 µg/ml ampicillin and 10 µg/ml tetracycline. Expression of GFP-FtsI was confirmed in the transductant clones by induction with 10-20 µM IPTG. SD247 (W3110 *ftsW*^*M269I*^) was previously described (12) and PK247-4 [SD247 *ftsL::kan/*pSD296] was generated by P1 transduction of *ftsL::kan* from the SD399 donor to the recipient strain SD247/pSD296 [P_ara_::*ftsL*]) and selecting Kan resistance and screening for arabinose dependency. PK4-1 (*ftsL::kan/*pKTP108 [P_ara_::*ftsL*]) was generated by using the same procedure described above. Unless stated otherwise, Luria-Broth (LB) medium containing 0.5% NaCl at indicated temperatures was used. For selection on LB agar and growth in LB broth, the following antibiotics and reagents were added at the indicated final concentrations as necessary (ampicillin, 100 μg/ml; spectinomycin, 50 μg/ml; kanamycin, 25 µg/ml; chloramphenicol, 10 µg/ml; tetracycline, 10 µg/ml; IPTG. 10-200 µM; glucose, 0.2%; and arabinose, 0.2%.

### Plasmids

Plasmids are listed in Table S1B. Genomic DNA extracted from W3110 strain was used as a template to obtain PCR fragments togenerate expression plasmids for *ftsL*. To construct the following plasmids: pKTP100 (P_*tac*_::*ftsL*), pKTP102 (P_*tac*_::*torA*^*1-42*^ *ftsL*^*58-121*^*-6xhis*), and pKTP103 [P_*tac*_::*malF*^*1-37*^ *ftsL*^*58-121*^*-6xhis*], the *ftsL* ORF was PCR amplified using a strong ribosome binding site in the forward primers targeting *ftsL* that included sequences encoding *torA*^*1-42*^ and *malF*^*1-37*^, respectively. The PCR fragments were digested with *Eco*RI and *Hin*dIII and ligated into the same sites in pJF118EH. Construction of pKTP104 (*P*_*T5*_::*ftsL*) and pKTP105 (P_T5_::*ftsL*^*30-121*^) involved PCR amplification of the *ftsL* ORF, digestion with *Bam*HI and *Hin*dIII, and ligation into the same sites in the pQE80L vector (Qiagen). The construction of pKTP108 [*repA*^ts^ P_syn135_::*ftsL*) employed a similar approach to that used for of pSD256 (12) except a strong ribosome binding site was added and the *Xba*I site was used instead of *Eco*RI. To create pKTP109, the *ftsI* ORF was PCR amplified and digested with SacI and HindIII followed by ligation into pBAD33 using sites with compatible overhangs. To generate plasmid pSD296 (P_ara_::*ftsL*), the *ftsL* ORF and its flanking sequences (250 bp) were PCR-amplified, digested with *Xba*I and *Hin*dIII, and ligated into the same sites in the pBAD33 vector. Plasmids pKTP106 (P_ara_::*ftsL*) and pKTP107 (P_ara_::*ftsL*^*30-121*^) were created by PCR amplification of *ftsL* and *ftsL*^*30-121*^, respectively using the primers that contain the same ribosome binding site as in pKTP100. The two PCR fragments were cut with *Sac*I and *Hin*dIII and cloned into sites in pBAD33 with compatible overhangs. The pND16 [P_*ftsK*_::*ftsW-ftsK*^*179-1329*^] plasmid constitutively expresses the FtsW-FtsK C-terminal fusion protein and pBL154 (repA^TS^ P_syn135_::*ftsN*) were previously described (32, 33). The overexpression plasmids for FtsN and FtsW, pSEB417 (P_204_::*ftsN*) and pSEB429 (P_204_:: *ftsW)*, respectively, were previously described (21, 32). Bacterial two hybrid (BTH) vectors, pUT18 (Amp^R^, cyaAT18 fragment) and pKT25 (Kan^R^, cyaAT25 fragment), were described previously (34). pUT18-ftsL (*cya*^*T18*^*-ftsL*) and pUT18-ftsL^30-121^ (*cya*^*T18*^*-ftsL*^*30-121*^) plasmids were generated by ligating PCR-amplified *ftsL* and *ftsL*^*30-121*^ into pUT18 (*cya*^*T18*^) digested with *Bam*HI *and Eco*RI, respectively. Construction of pUT18-ftsW (*cya*^*T18*^*-ftsW*) and pKT25-ftsW (*cya*^*T25*^*-ftsW*) involved PCR amplification of *E. coli ftsW ORF* and digestion of the fragments with *Bam*HI and *Kpn*I followed by ligation into the BTH vectors digested with the same enzymes. pKT25-ftsI (*cya*^*T25*^*-ftsI*) was created by similar procedures but *Bam*HI and *Eco*RI were used for digestion of PCR fragment and vector. All primers are listed in Table S2.

### Random and Site-Directed Mutagenesis

To obtain the *ftsL* and *ftsB* mutant libraries (with a single missense mutation per *ORF*) an optimal mutation rate (0.3–1 base/kb) for 1µg of template was adopted as recommended in GeneMorph II Random Mutagenesis kit (Agilent Technologies). The PCR products were then digested with *Eco*RI and *Hin*dIII and ligated into the pJF118EH vector using the same restriction enzymes. A ligation pool of pJF118EH-ftsL or pJF118EH-ftsB containing putative mutations was transformed into JS238 by electroporation and a library was prepared by isolating plasmids from a pool of growing colonies. A dominant negative phenotype, largely characterized by flat colony morphology was screened for by introducing the resulting plasmids into JS238. Specific point mutations in *ftsL, ftsL*^*30-121*^ and *ftsW* were introduced into some plasmids by using the QuickChange site-directed mutagenesis kit according to the manufacturer’s instruction (Agilent Technologies).

### Isolation of an allele of *ftsW* that bypasses *ftsN*

To generate a library of random *ftsW* mutations, *ftsW* was subjected to random PCR mutagenesis and cloned into plasmid pSEB429 (P_204_::*ftsW*) to replace the WT *ftsW*. The mutagenized library (pSEB429M) was transformed into strain SD399 [*ftsL::kan* /pSD256 (repA^ts^::*ftsL)*] harboring plasmid pSD296-E87K (P_ara_::*ftsL*^*E87K*^) and suppressors of FtsL^E87K^ were selected on LB plates with 0.2% arabinose (to induce *ftsL*^*E87K*^) and 60 µM IPTG (to induce *ftsW*) at 37 °C. Fourteen of the surviving clones were purified, retested and sequenced. Eleven contained a single mutation (E289G) while 3 contained this mutation plus other mutations. The *ftsW*^*E289G*^ mutation was introduced into S3 (W3110, *leu::Tn10*) by recombineering. P1 transduction of *ftsN::kan* from strain CH34/pMG20 (*ftsN::kan*) into SD488 (*leu::Tn10, ftsW*^*E289G*^) was done using following a standard procedure. The Kan^R^ transductants had a slightly longer phenotype than a WT strain.

### Helix Modeling of FtsL periplasmic domain

A secondary structure of FtsL was generated for illustrative purposes. To do this, a crude model of the putative coiled coil region of FtsL was modelled on the coiled coil structure (tropomyosin, 1IC2). Structures were visualized using PyMOL (Molecular Graphics System Version 1.2r3pre, Schrödinger, LLC).

### Bacterial Two-Hybrid Analysis

The *cya* null strain DHM1 [*F-, cya-854, recA1, endA1, gyrA96 (Nal*^*r*^*), thi1, hsdR17, spoT1, rfbD1, glnV44(AS)*] was simultaneously transformed with plasmids pKT25-ftsW or pKT25-ftsI and pUT18-ftsL (or-ftsL^30-121^), carrying wild-type or mutant *ftsW* and *ftsL* alleles and grown overnight at 30°C on LB plates containing 0.2% glucose, 25 μg/ml kanamycin, and 100 μg/ml ampicillin. Colonies from the LB plates were diluted in 300 μl volume of LB broth and spotted onto fresh LB plates supplemented with 25 μg/ml kanamycin, 100 μg/ml ampicillin, 40 μg/ml 5-bromo-4-chloro-3-indoyl-β-D-galactopyranoside (X-Gal), and 0.5 mM IPTG. The color changes were recorded after overnight incubation at room temperature at 30°C.

### Microscopy

The dominant negative effects of the FtsL mutants on cell division were assessed using phase-contrast microscopy by monitoring the degree of filamentation. JS238 containing pKTP100 or derivatives carrying *ftsL* mutations was grown overnight in the presence of 100 μg/ml ampicillin and 0.2% glucose and the cultures were diluted 1/200-1/500 in fresh LB medium containing 100 μg/ml ampicillin. At OD_540_ ≈ 0.02, 50 µM IPTG was added and cell morphologies were analyzed 2 hours later.

To visualize GFP-FtsI localization, SD285 [*leu::*Tn*10 bla lacI*^q^ *P*_207_-*gfp-ftsI*] containing pKTP106 (Para::*ftsL*) or derivatives with the *ftsL*^*E87K*^ or *ftsL*^*A90E*^ mutations was grown overnight at 30°C in LB medium containing 50 μg/ml ampicillin and 10 μg/ml chloramphenicol. The overnight cultures were diluted 1/200 ∼ 1/500 in fresh LB medium containing the same antibiotics, 0.2% arabinose, and 10-20 µM IPTG, and were incubated at 37°C until OD_540_≈ 0.4. Cells were immobilized on an LB agarose pad and the localization of GFP-FtsI was recorded using a cooled CCD camera and processed using Metamorph (Molecular Devices) and Adobe Photoshop.

## Acknowledgements

We would like to thank Piet de Boer and David Weiss for strains and plasmids and Scott Lovell for generating the model of the FtsL alpha helix. This study was supported by NIH grant GM29746 to Joe Lutkenhaus.

## Author contributions

K.P. and J.L. designed the research; K.P. and S.D. performed the research; K.P., S.D. and J.L. analyzed data and wrote the manuscript.

## Notes

### Competing Interest Statement

The authors have declared no competing interest.

## References

1. de Boer PA (2010) Advances in understanding E. coli cell fission. Curr Opin Microbiol 13(6):730–737.

2. Du S & Lutkenhaus J (2017) Assembly and activation of the Escherichia coli divisome. Mol Microbiol 105(2):177–187.

3. Goehring NW, Gonzalez MD, & Beckwith J (2006) Premature targeting of cell division proteins to midcell reveals hierarchies of protein interactions involved in divisome assembly. Mol Microbiol 61(1):33–45.

4. Yang X, et al. (2017) GTPase activity-coupled treadmilling of the bacterial tubulin FtsZ organizes septal cell wall synthesis. Science 355(6326):744–747.

5. Spratt BG (1975) Distinct penicillin binding proteins involved in the division, elongation, and shape of Escherichia coli K12. Proc Natl Acad Sci U S A 72(8):2999–3003.

6. Meeske AJ, et al. (2016) SEDS proteins are a widespread family of bacterial cell wall polymerases. Nature 537(7622):634–638.

7. Cho H, et al. (2016) Bacterial cell wall biogenesis is mediated by SEDS and PBP polymerase families functioning semi-autonomously. Nat Microbiol:16172.

8. Taguchi A, et al. (2019) FtsW is a peptidoglycan polymerase that is functional only in complex with its cognate penicillin-binding protein. Nat Microbiol.

9. Addinall SG, Cao C, & Lutkenhaus J (1997) FtsN, a late recruit to the septum in Escherichia coli. Mol Microbiol 25(2):303–309.

10. Liu B, Persons L, Lee L, & de Boer PA (2015) Roles for both FtsA and the FtsBLQ subcomplex in FtsN-stimulated cell constriction in Escherichia coli. Mol Microbiol 95(6):945–970.

11. Tsang MJ & Bernhardt TG (2015) A role for the FtsQLB complex in cytokinetic ring activation revealed by an ftsL allele that accelerates division. Mol Microbiol 95(6):925–944.

12. Du S, Pichoff S, & Lutkenhaus J (2016) FtsEX acts on FtsA to regulate divisome assembly and activity. Proc Natl Acad Sci U S A 113(34):E5052–5061.

13. Gonzalez MD, Akbay EA, Boyd D, & Beckwith J (2010) Multiple interaction domains in FtsL, a protein component of the widely conserved bacterial FtsLBQ cell division complex. J Bacteriol 192(11):2757–2768.

14. Robichon C, Karimova G, Beckwith J, & Ladant D (2011) Role of leucine zipper motifs in association of the Escherichia coli cell division proteins FtsL and FtsB. J Bacteriol 193(18):4988–4992.

15. Condon SGF, et al. (2018) The FtsLB subcomplex of the bacterial divisome is a tetramer with an uninterrupted FtsL helix linking the transmembrane and periplasmic regions. J Biol Chem 293(5):1623–1641.

16. Khadria AS & Senes A (2013) The transmembrane domains of the bacterial cell division proteins FtsB and FtsL form a stable high-order oligomer. Biochemistry 52(43):7542–7550.

17. Buddelmeijer N & Beckwith J (2004) A complex of the Escherichia coli cell division proteins FtsL, FtsB and FtsQ forms independently of its localization to the septal region. Mol Microbiol 52(5):1315–1327.

18. Gonzalez MD & Beckwith J (2009) Divisome under construction: distinct domains of the small membrane protein FtsB are necessary for interaction with multiple cell division proteins. J Bacteriol 191(8):2815–2825.

19. Kureisaite-Ciziene D, et al. (2018) Structural Analysis of the Interaction between the Bacterial Cell Division Proteins FtsQ and FtsB. MBio 9(5).

20. Choi Y, et al. (2018) Structural Insights into the FtsQ/FtsB/FtsL Complex, a Key Component of the Divisome. Sci Rep 8(1):18061.

21. Pichoff S, Du S, & Lutkenhaus J (2015) The bypass of ZipA by overexpression of FtsN requires a previously unknown conserved FtsN motif essential for FtsA-FtsN interaction supporting a model in which FtsA monomers recruit late cell division proteins to the Z ring. Mol Microbiol 95(6):971–987.

22. Busiek KK, Eraso JM, Wang Y, & Margolin W (2012) The early divisome protein FtsA interacts directly through its 1c subdomain with the cytoplasmic domain of the late divisome protein FtsN. J Bacteriol 194(8):1989–2000.

23. Mercer KL & Weiss DS (2002) The Escherichia coli cell division protein FtsW is required to recruit its cognate transpeptidase, FtsI (PBP3), to the division site. J Bacteriol 184(4):904–912.

24. Sjodt M, et al. (2020) Structural coordination of polymerization and crosslinking by a SEDS-bPBP peptidoglycan synthase complex. Nat Microbiol 5(6):813–820.

25. Wissel MC & Weiss DS (2004) Genetic analysis of the cell division protein FtsI (PBP3): amino acid substitutions that impair septal localization of FtsI and recruitment of FtsN. J Bacteriol 186(2):490–502.

26. Guzman LM, Weiss DS, & Beckwith J (1997) Domain-swapping analysis of FtsI, FtsL, and FtsQ, bitopic membrane proteins essential for cell division in Escherichia coli. J Bacteriol 179(16):5094–5103.

27. Dubarry N & Barre FX (2010) Fully efficient chromosome dimer resolution in Escherichia coli cells lacking the integral membrane domain of FtsK. EMBO J 29(3):597–605.

28. LaPointe LM, et al. (2013) Structural organization of FtsB, a transmembrane protein of the bacterial divisome. Biochemistry 52(15):2574–2585.

29. Boes A, Olatunji S, Breukink E, & Terrak M (2019) Regulation of the Peptidoglycan Polymerase Activity of PBP1b by Antagonist Actions of the Core Divisome Proteins FtsBLQ and FtsN. mBio 10(1).

30. Contreras-Martel C, et al. (2017) Molecular architecture of the PBP2-MreC core bacterial cell wall synthesis complex. Nat Commun 8(1):776.

31. Rohs PDA, et al. (2018) A central role for PBP2 in the activation of peptidoglycan polymerization by the bacterial cell elongation machinery. PLoS Genet 14(10):e1007726.

32. Dubarry N, Possoz C, & Barre FX (2010) Multiple regions along the Escherichia coli FtsK protein are implicated in cell division. Mol Microbiol 78(5):1088–1100.

33. Gerding MA, et al. (2009) Self-enhanced accumulation of FtsN at Division Sites and Roles for Other Proteins with a SPOR domain (DamX, DedD, and RlpA) in Escherichia coli cell constriction. J Bacteriol 191(24):7383–7401.

34. Karimova G, Pidoux J, Ullmann A, & Ladant D (1998) A bacterial two-hybrid system based on a reconstituted signal transduction pathway. Proc Natl Acad Sci U S A 95(10):5752–5756.

