## Supplemental figures and tables for "Essential role for FtsL in activation of septal PG synthesis": Supplementary Material; Park et al.pdf

By

Kyung-Tae Park, Shishen Du and Joe Lutkenhaus

Contents:

Figures – Fig. S1-S9 with legends

Tables – Table S1-S2

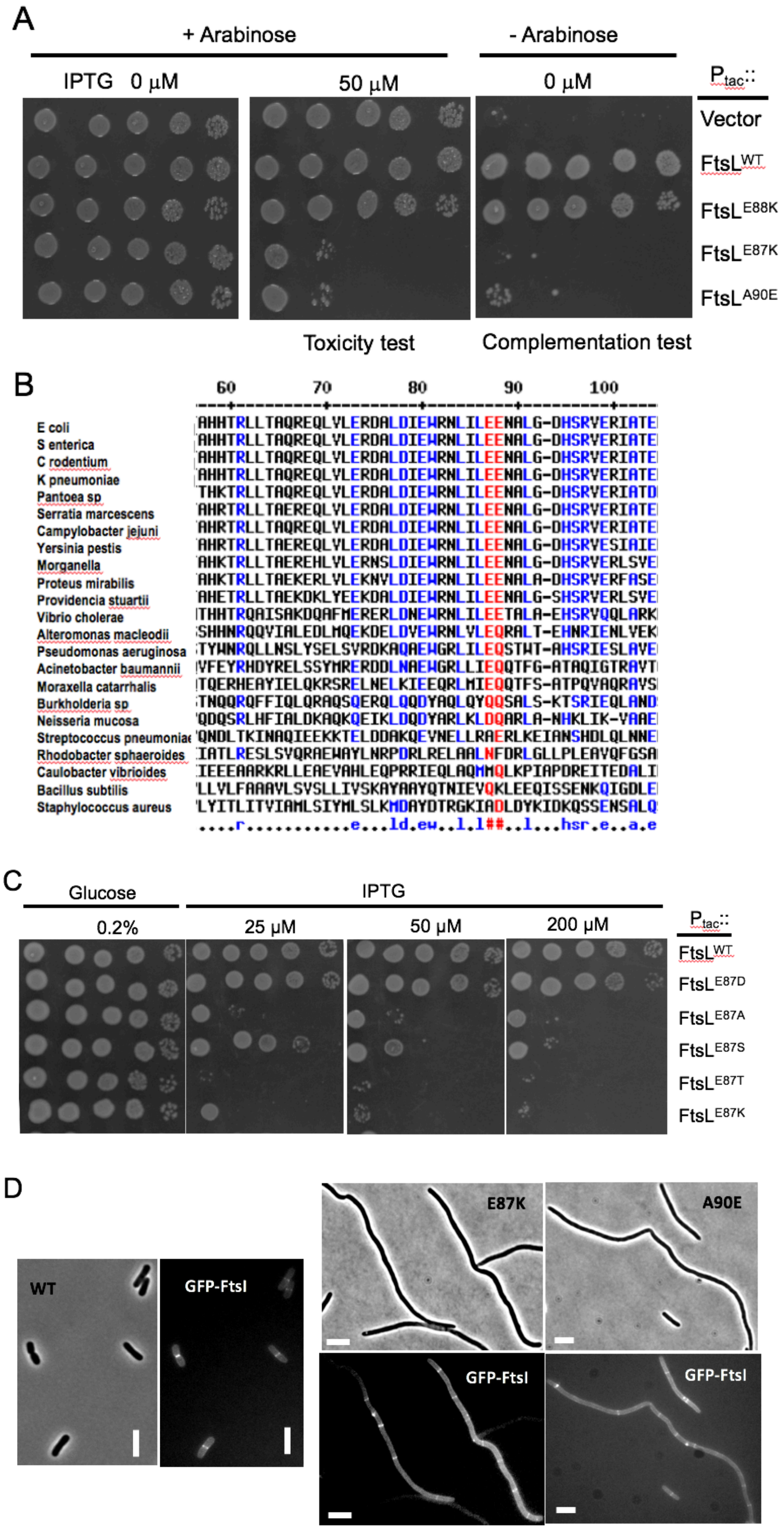
A
+ Arabinose
- Arabinose
IPTG 0 μM
50 μM
0 μM
Ptac::
Vector
FtsL WT
FtsL E88K
FtsL E87K
FtsL A90E
Toxicity test
Complementation test
B
60
70
80
90
100
E coli
S enterica
C rodentium
K pneumoniae
Pantoea sp
Serratia marcescens
Campylobacter jejuni
Yersinia pestis
Morganella
Proteus mirabilis
Providencia stuartii
Vibrio cholerae
Alteromonas macleodii
Pseudomonas aeruginosa
Acinetobacter baumannii
Moraxella catarrhalis
Burkholderia sp
Neisseria mucosa
Streptococcus pneumoniae
Rhodobacter sphaeroides
Caulobacter vibrioides
Bacillus subtilis
Staphylococcus aureus
C
Glucose
IPTG
0.2%
25 μM
50 μM
200 μM
Ptac::
FtsL WT
FtsL E87D
FtsL E87A
FtsL E87S
FtsL E87T
FtsL E87K
D
WT
GFP-FtsI
E87K
A90E
GFP-FtsI
GFP-FtsI

Fig. S1. Characterization of dominant negative mutations in *ftsL*. A) Dominant negative mutations do not complement *ΔftsL*. Two dominant negative alleles of *ftsL* were tested for their ability to complement an *ftsL* depletion strain by transforming SD439 (*ftsL::kan*) /pSD296 ($P_{ara}$::*ftsL*) with pKTP100 ($P_{tac}$::*ftsL*) derivatives carrying different *ftsL* alleles. The toxicity of these alleles was tested on plates containing both arabinose (induced WT *ftsL*) and IPTG (to induce the allele to be tested) (center panel). The ability to complement the *ftsL* depletion strain was assessed on plates without arabinose or IPTG (basal level of expression from pKTP100 ($P_{tac}$::*ftsL*) is sufficient for complementation) (right panel). B) Lineup of a region (amino acids 57-104) of the periplasmic domain of FtsL. FtsL from a variety of Gram-negative and Gram-positive bacteria were aligned using Cobalt. Residues in red are more conserved than residues in blue which are more conserved than those in gray. Only residues E87 and E88 are colored red. C) Examination of additional substitutions at position 87 in FtsL on toxicity. Derivatives of pKT100 ($P_{tac}$::*ftsL*) carrying various alleles of *ftsL* were introduced into JS238 and tested for toxicity by performing spot tests on plates containing increasing amounts of IPTG. D) Effect of the dominant negative *ftsL* mutations on the recruitment of GFP-FtsI. SD285 (*leu*::Tn*10 recA::aadA* $P_{206}$::*gfp-ftsI*) was transformed with pSD296 ($P_{ara}$::*ftsL*) derivatives expressing $ftsL^{E87K}$, $ftsL^{A90E}$ or *ftsL*. Two hours after induction with 0.2% arabinose samples were taken and examined by phase and fluorescent microscopy.

A

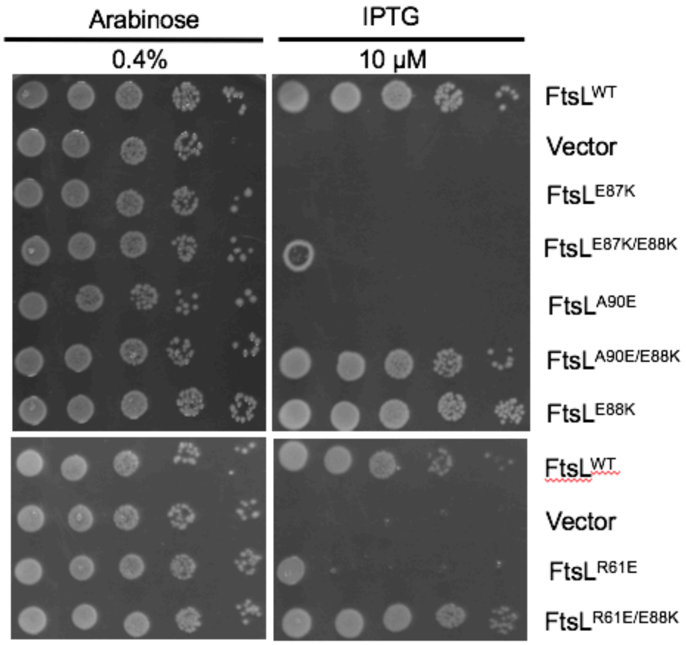

Arabinose
0.4%
IPTG
10 μM
FtsL WT
Vector
FtsL E87K
FtsL E87K/E88K
FtsL A90E
FtsL A90E/E88K
FtsL E88K
FtsL WT
Vector
FtsL R61E
FtsL R61E/E88K

B

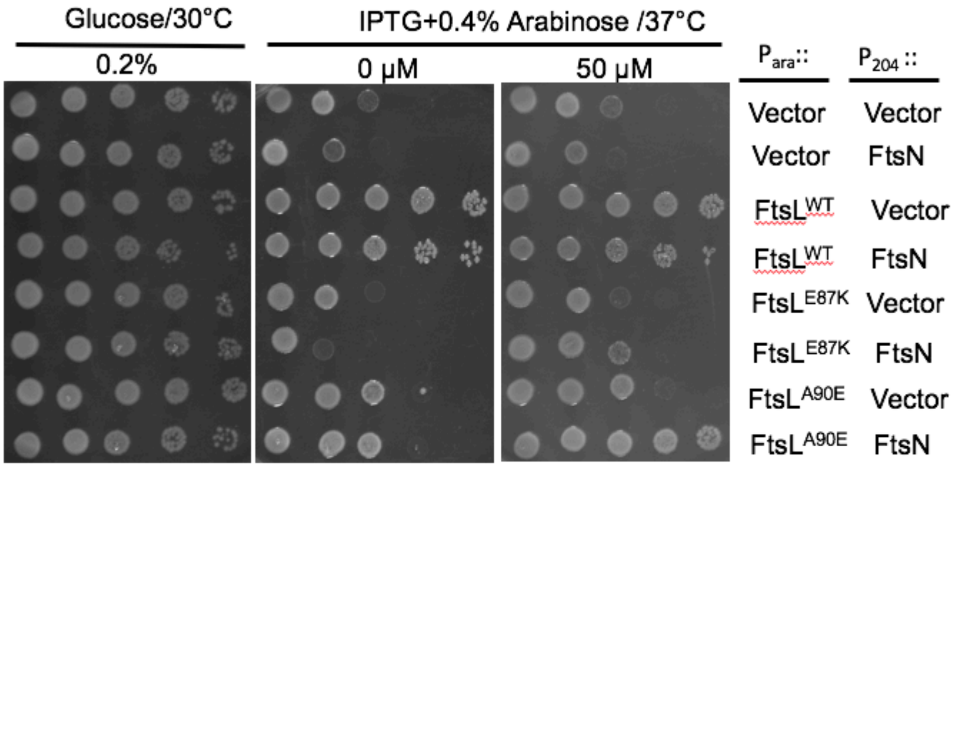

Glucose/30°C
0.2%
IPTG+0.4% Arabinose /37°C
0 μM
50 μM
P ara ::
P 204 ::
Vector Vector
Vector FtsN
FtsL WT Vector
FtsL WT FtsN
FtsL E87K Vector
FtsL E87K FtsN
FtsL A90E Vector
FtsL A90E FtsN

Fig. S2. Test for rescue of dominant negative *ftsL* mutations by an *ftsL* activation mutation or by *ftsN* overexpression. A) Rescue by an *ftsL* activation mutation in *cis*. The $ftsL^{E88K}$ mutation was added to various *ftsL* alleles in cis and then tested for complementation. To do this SD439 (*ftsL::kan*/pSD296 [$P_{ara}$::*ftsL*]) was transformed with derivatives of pKTP100 ($P_{tac}$::*ftsL*) carrying the various mutations. The strains were spotted on plates at 30°C without arabinose to deplete WT *ftsL* and IPTG added to induce the various *ftsL* alleles. $ftsL^{A90E}$ and $ftsL^{R61E}$ were rescued but $ftsL^{E87K}$ and $ftsL^{L86F}$ (Table 1) were not. B) Rescue by overexpression of *ftsN*. To test if *ftsN* overexpression can rescue any of the dominant negative alleles of *ftsL,* SD399 (*ftsL::kan*/pSD256 [$repA^{TS}$ $P_{syn135}$::*ftsL*]) containing plasmids expressing the *ftsL* alleles under an arabinose inducible promoter (derivatives of pSD296 [$P_{ara}$::*ftsL*]) were transformed with a plasmid with *ftsN* under IPTG control (pSEB417 [$P_{204}$::*ftsN*]). The strains were spotted at 37°C to deplete WT *ftsL* and arabinose added to induce the *ftsL* allele and IPTG added to induce *ftsN*.

A

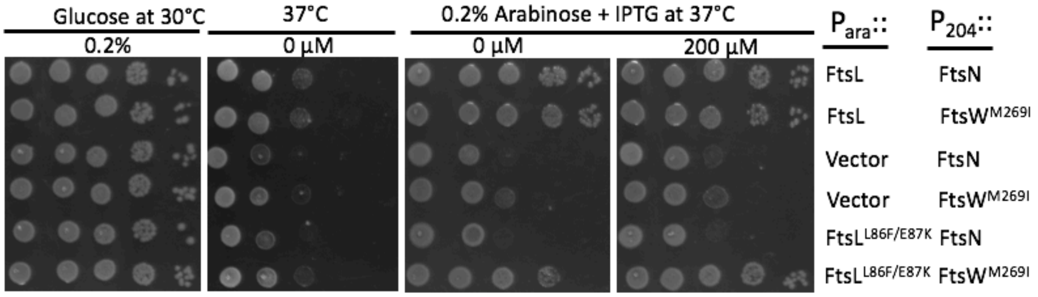

B

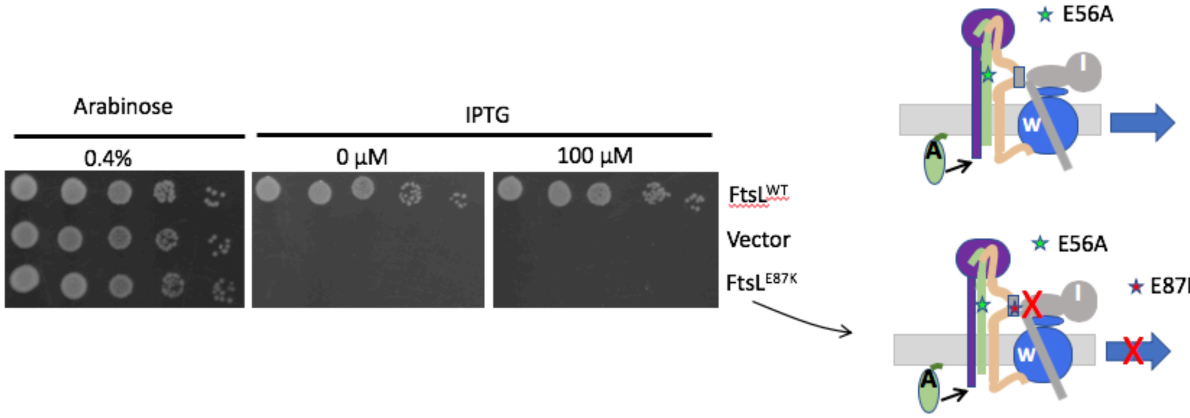

Fig. S3. Effect of activation mutations in *ftsW* and *ftsB* and *ftsN* overexpression on the rescue of dominant negative *ftsL* mutations. A) Overexpression of an activation allele of *ftsW*, but not *ftsN*, suppresses a dominant negative allele of *ftsL (ftsL*$^{L86F/E87K}$*)*. SD399 (*ftsL::kan*/pSD256[*repA*$^{ts}$-*ftsL*]) was transformed with plasmids expressing either *ftsL* or a dominant negative allele of *ftsL* (*ftsL*$^{E87K/L86F}$) under an arabinose-inducible promoter (derivatives of pSD296 [P$_{ara}$::*ftsL*]) and plasmids expressing *ftsW*$^{M269I}$ or *ftsN* under an IPTG-inducible promoter (pSEB429-I [P$_{204}$::*ftsW*$^{M269I}$] and pSEB417 [P$_{204}$::*ftsN*], respectively). The strains were spotted on plates at 37°C (to deplete WT *ftsL*) containing 0.2% arabinose to induce the *ftsL* alleles and increasing concentrations of IPTG to induce *ftsW*$^{M269I}$ or *ftsN*. The second panel lacks arabinose demonstrating that the *ftsW* activation mutations cannot bypass *ftsL.* B)

An activated allele of *ftsB* cannot suppress a dominant negative *ftsL* mutation. Strain BL167 (*ftsB*$^{E56A}$ *recA::aadA ftsL::kan*/pSD296 [$P_{ara}$::*ftsL*]) containing pKTP100 ($P_{tac}$::*ftsL*) or a version expressing *ftsL*$^{E87K}$ was subjected to spot tests on plates without arabinose to deplete WT *ftsL* and with IPTG added to induce the *ftsL* allele cloned in derivatives of pKTP100.

A

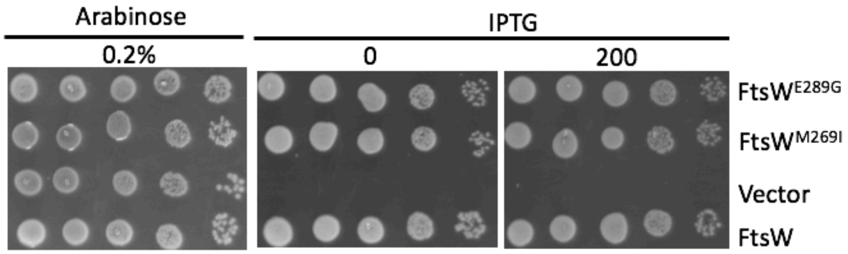

B

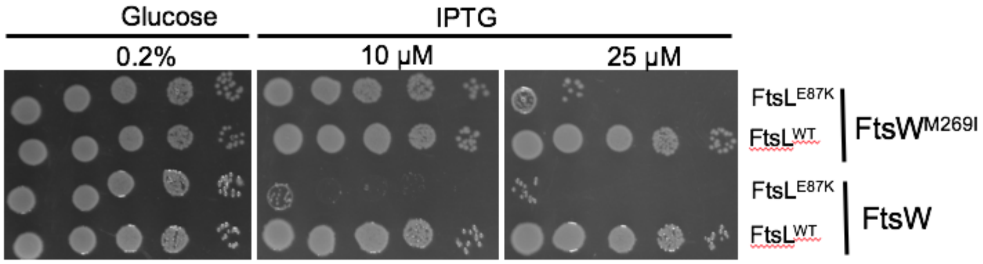

Fig. S4. Effect of activation mutations in *ftsW* on complementation and rescue of a dominant negative *ftsL* allele. A) Test of *ftsW* alleles for complementation of a *ftsW* depletion strain in the absence of ITPG. To test if the addition of IPTG was required for complementation of an *ftsW* depletion strain, EC912 (*ftsW::kan*/pDSW406 [$P_{ara}$:: *ftsW*]) was transformed with derivatives of pSEB429 ($P_{204}$::*ftsW*) carrying various alleles of *ftsW*. The strains were spot tested on plates without arabinose and with or without 200 μM IPTG. B) Expression of *ftsL*$^{E87K}$ inhibits a strain with the *ftsW*$^{M269I}$ mutation on the chromosome. Strains KTP1 (*ftsW*$^{+}$) and SD247-1 (*ftsW*$^{M269I}$) were transformed with pKTP100 ($P_{tac}$::*ftsL*) or a derivative expressing *ftsL*$^{E87K}$. The strains were spot tested on plates containing increasing concentrations of IPTG to test for toxicity. The strain containing the *ftsW*$^{M269I}$ is more resistant but still sensitive to *ftsL*$^{E87K}$.

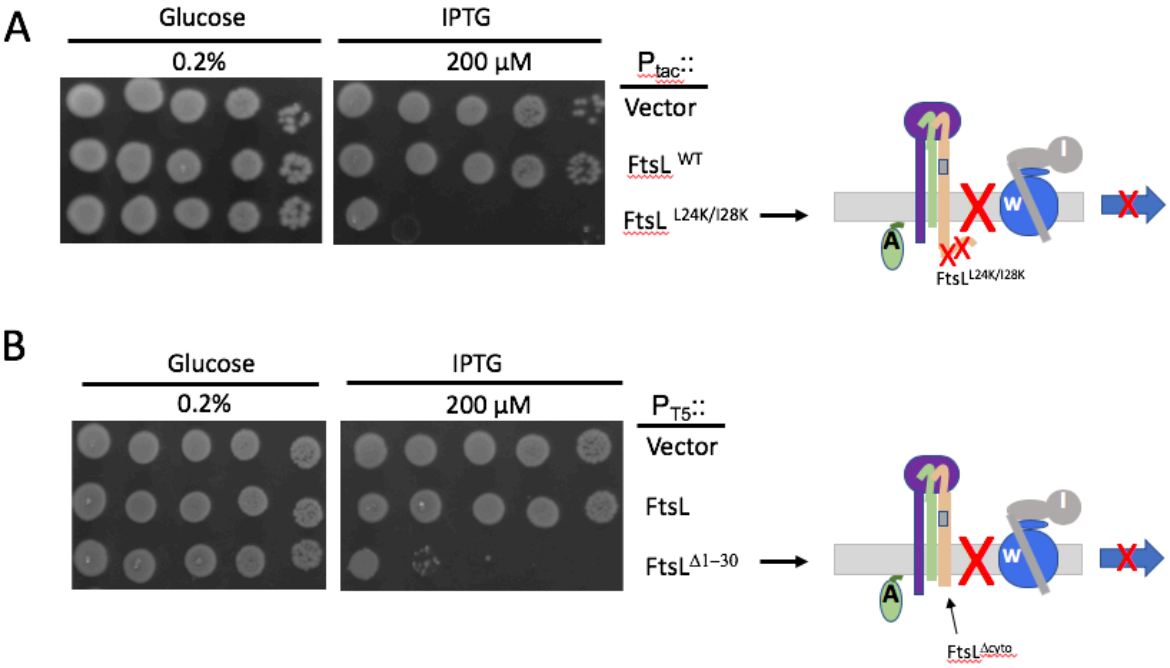

Fig. S5. Mutations in the cytoplasmic domain of *ftsL* result in a dominant negative phenotype. A) Mutations (L24K, I28K) in the ${}^{cyto}$*ftsL* domain lead to a dominant negative phenotype. JS238 containing pKTP100 ($P_{tac}$::*ftsL*) derivatives expressing different alleles of *ftsL* were tested for toxicity following induction with IPTG. B) Deletion of the cytoplasmic domain of FtsL (*ftsL*$^{\Delta 1\text{-}30}$) results in a dominant negative allele. JS238 containing pKTP104 ($P_{T5}$::*ftsL*) or pKTP105 ($P_{T5}$::*ftsL*$^{\Delta 1\text{-}30}$) were tested for toxicity following induction with IPTG.

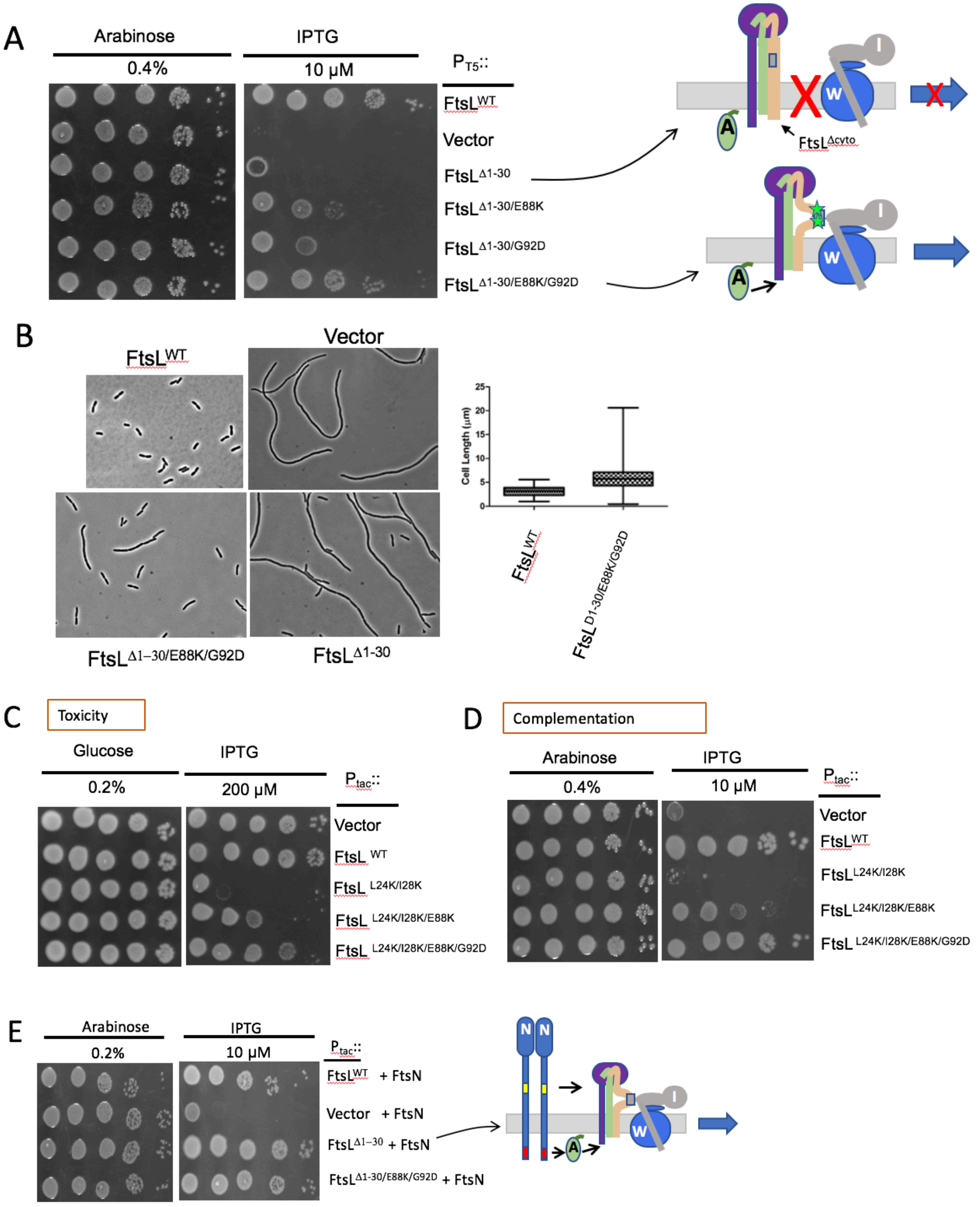
A
Arabinose
0.4%
IPTG
10 µM
PT5::
FtsLWT
Vector
FtsLΔ1-30
FtsLΔ1-30/E88K
FtsLΔ1-30/G92D
FtsLΔ1-30/E88K/G92D
FtsLΔcyto
A
W
I
B
FtsLWT
Vector
FtsLΔ1–30/E88K/G92D
FtsLΔ1-30
Cell Length (µm)
25
20
15
10
5
0
FtsL WT
FtsL D1-30/E88K/G92D
C
Toxicity
Glucose
0.2%
IPTG
200 µM
Ptac::
Vector
FtsL WT
FtsL L24K/I28K
FtsL L24K/I28K/E88K
FtsL L24K/I28K/E88K/G92D
D
Complementation
Arabinose
0.4%
IPTG
10 µM
Ptac::
Vector
FtsLWT
FtsLL24K/I28K
FtsLL24K/I28K/E88K
FtsL L24K/I28K/E88K/G92D
E
Arabinose
0.2%
IPTG
10 µM
Ptac::
FtsLWT + FtsN
Vector + FtsN
FtsLΔ1–30 + FtsN
FtsLΔ1-30/E88K/G92D + FtsN
N
N

Fig. S6. Rescue of *ftsL* cytoplasmic mutations by *ftsL* activation mutations and *ftsN* overexpression. A**)** *ftsL*$^{\Delta 1\text{-}30}$ rescue require two *ftsL* activation mutations. SD439 (*ftsL::kan*/pSD296 [$P_{ara}$::*ftsL*]) carrying pKTP105 ($P_{T5}$::*ftsL*$^{\Delta 1\text{-}30}$) derivatives expressing various alleles of *ftsL* were tested for their ability to complement an *ftsL* depletion strain. Complementation was tested by spotting strains on plates without arabinose (to deplete WT *ftsL*) and with ITPG to induce the various *ftsL* alleles. Loss of a functional $^{cyto}$FtsL domain (and therefore the ability to recruit FtsW) is compensated for by the two activation mutations *ftsL*$^{E88K/G92D}$. Cartoon depicting the rescue by the activation mutations. B**)** Morphology of *ΔftsL* cells rescued by *ftsL*$^{\Delta 1\text{-}30}$ carrying activation mutations. SD339 (*ftsL::kan*/pSD256 [$P_{ara}$::*ftsL*]) carrying derivatives of pKTP107 [$P_{ara}$::*ftsL*] expressing various alleles of *ftsL*$^{\Delta 1\text{-}30}$ (from Panel A) were grown to exponential phase at 30$^{o}$C with antibiotics, centrifuged, washed and resuspended in LB with 0.2% arabinose and antibiotics at 30$^{o}$C. Samples were taken 2.5 hours later for photography and the cell length distributions determined. C) *ftsL* activation mutations reduce the toxicity of *ftsL*$^{L24K/I28K}$. JS238 carrying derivatives of pKTP100 ($P_{tac}$::*ftsL*) expressing different alleles of *ftsL* was tested for toxicity by spotting on plates containing 200 μM IPTG to induce the *ftsL* alleles. D) *ftsL* activation mutations rescue *ftsL*$^{L24K/I28K}$ for complementation. SD439 (*ftsL::kan*/pSD296 [$P_{ara}$::*ftsL*]) containing derivatives of pKTP100 [$P_{tac}$::*ftsL*]) with various *ftsL* alleles was spotted on plates without arabinose (to deplete WT *ftsL*) and with IPTG to induce the mutant *ftsL* alleles. E) Overexpression of *ftsN* suppresses *ftsL*$^{\Delta 1\text{-}30}$. SD439 (*ftsL::kan*/pSD296 [$P_{ara}$::*ftsL]*) containing derivatives of pKTP105 ($P_{T5}$::*ftsL* $^{\Delta 1\text{-}30}$) with various alleles of *ftsL* was transformed with pBL154 (*repA*$^{TS}$ $P_{ftsN}$::*ftsN*) which constitutively expresses *ftsN*. Rescue of FtsL$^{\Delta 1\text{-}30}$ was tested by spotting transformants on plates without arabinose and with IPTG.

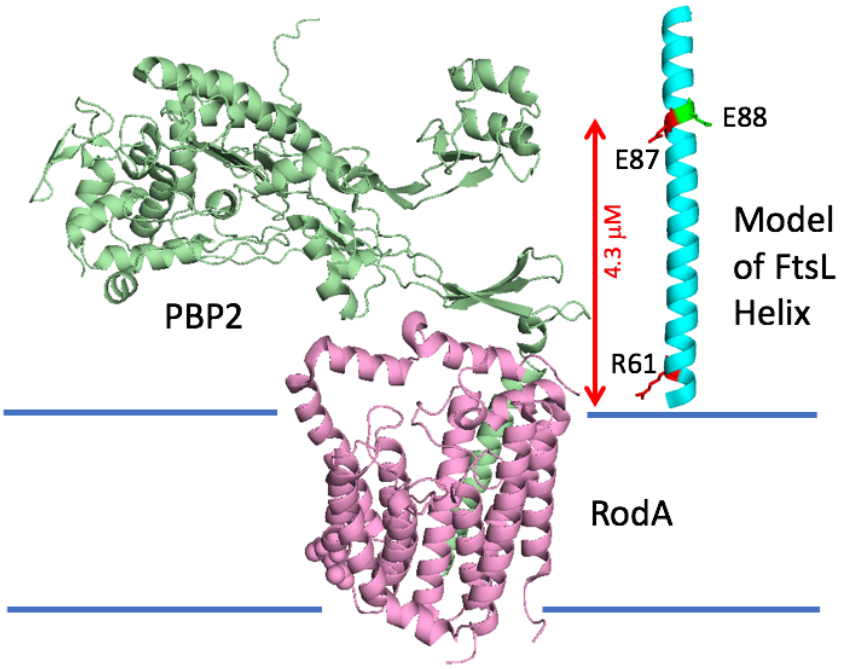

Fig. S7. Diagram indicating the position AWI of FtsL relative to the membrane and the RodA-PBP2 complex. Part of the periplasmic domain of FtsL was modeled as an alpha helix (residues 57-99) and positioned next to the structure of the RodA-PBP2 complex (PDB ID: 6PL6). The RodA-PBP2 complex is homologous to the FtsW-PBP3 (FtsI) complex. Representative FtsL residues E87 (AWI), E88 (CCD) are about 4.3 μM from the membrane. The position of R61 is also indicated.

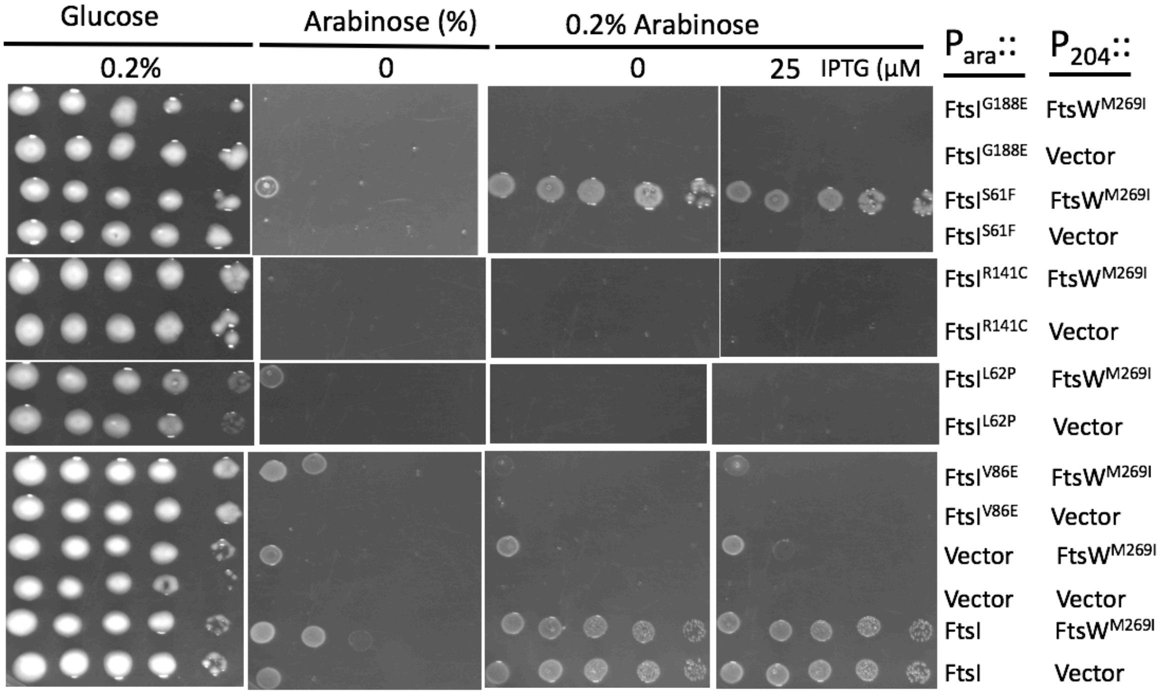

Fig. S8. Testing the rescue of additional alleles of *ftsI* by an activated FtsW mutant. Additional *ftsI* alleles were tested to see if they were rescued by *ftsW*$^{M269I}$ expression. Some alleles were tested in Fig. 6A and the others are tested here. Note among the alleles tested here only *ftsI*$^{S61F}$ is rescued. Also, the bottom row of panels is also presented in Fig. 6A and was included here for comparison. MCI23 (*ftsI23*$^{ts}$ *recA::spc)* was transformed with compatible plasmids expressing an activated allele of *ftsW* (pSEB429[$P_{204}$::*ftsW*$^{M269I}$]) and the various *ftsI* alleles under arabinose promoter control (derivatives of pBAD33-*ftsI*). Transformants were spot tested on plates at 37$^{o}$C (to inactivate *ftsI23*$^{Ts}$) and arabinose added to induce the *ftsI* alleles and increasing concentrations of IPTG to induce *ftsW*$^{M269I}$.

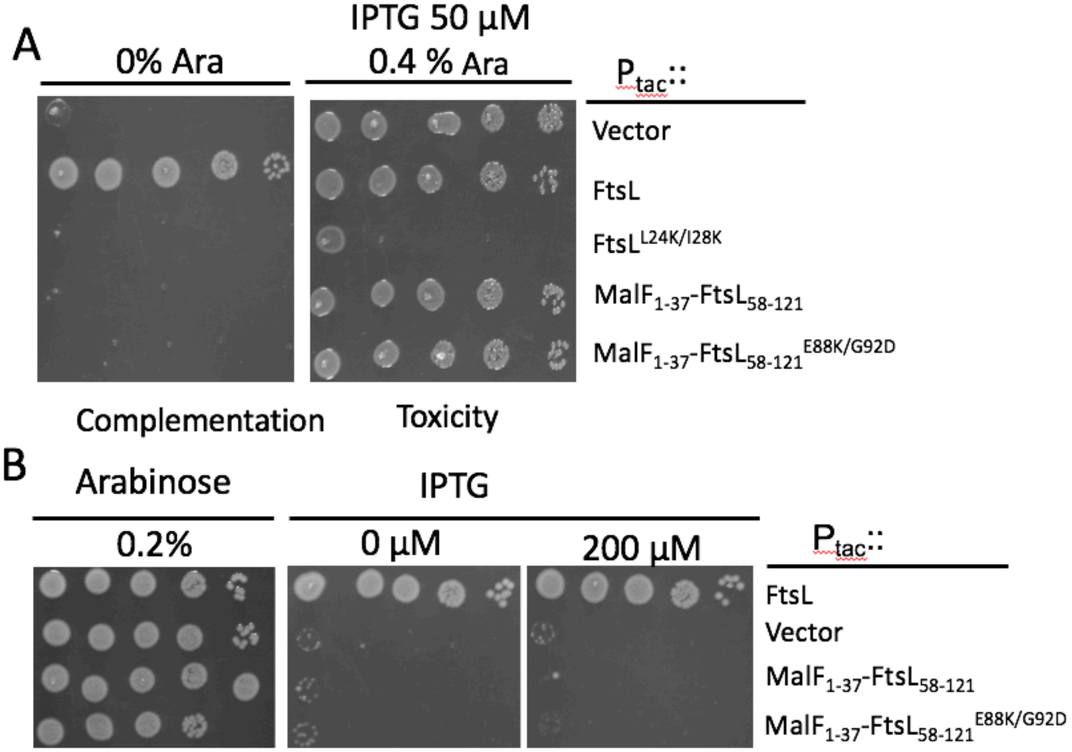

Fig. S9. Characterization of FtsL fusions. A) The MalF-FtsL fusion lacks toxicity and fails to complement a *ftsL* depletion strain. The left panel tests for complementation and the right panel tests for toxicity. SD439 (*ftsL::kan*/pSD296 [$P_{ara}$::*ftsL*]) containing pKTP100 ($P_{tac}$::*ftsL*) derivatives expressing various alleles of *ftsL* were incubated without arabinose or IPTG (basal expression from pKTP100 [$P_{tac}$::*ftsL*] is sufficient for complementation) to test for complementation. In the right panel both arabinose and IPTG were added. Arabinose induces WT *ftsL* from pSD296 [$P_{ara}$::*ftsL*] whereas IPTG induces the *ftsL* allele from pKTP100 ($P_{tac}$::*ftsL*). The $ftsL^{L24K,I28K}$ allele was added as a control to demonstrate toxicity. The lack of toxicity indicates that the MalF-FtsL fusion fails to form a complex with FtsQB. B) FtsL fusions are unable to complement a *ftsL* depleted strain. Fusion of the periplasmic domain of FtsL to the MalF cytoplasmic and transmembrane domains (with or without *ftsL* activation mutations) were tested for ability to complement an *ftsL* depletion strain. To do this PK247-4 ($ftsW^{M269I}$

*ftsL::kan*/pSD296 [$P_{ara}$::*ftsL*]) containing derivatives of pKTP100 ($P_{tac}$::*ftsL*) carrying various fusions in place of *ftsL* were spot tested on plates with IPTG.

Table S1. Strains and plasmids used in this study.

A. Strains

| Strains | Genotype | Source/ reference |
|---|---|---|
| AM1992 | *ΔrecA1921::spc* | E. coli Stock center |
| BL156/pJH2 | TB28, *ftsL::kan*/pJH2 ($P_{syn135}$::*ftsL*) | (1) |
| CH34/pMG20 | *ftsN::kan* | (1) |
| DHM1 | Bacterial 2-hybrid reporter strain MG1655 *F-, cya-854, recA1, endA1, gyrA96 (Nal r), thi1, hsdR17, spoT1, rfbD1, glnV44(AS)* | (2) |
| EC436 | *MC4100 Δ* (λ*attL-lom*)::*bla lacI*$^{q}$ *P*$_{207}$-*gfp-ftsI* | (3) |
| JS238 | MC1061 *malPp*::*lacI*$^{Q}$ *srlC*::Tn10 *recA1* | (4) |
| KTP1 | W3110 *recA::spc* | P1 on AM1992 X W3110, select Spc$^{R}$ |
| MCI23 | MC4100 *ftsI23* (Ts) | (5) |
| MCI23 recA | MC4100 *ftsI23* (Ts) *recA::spc* | P1 on AM1992 X MCI23, select Spc$^{R}$ |
| PK247-4 | SD247 *ftsL::kan/*pSD296 | P1 on SD399 X SD247, select kan$^{R}$ |
| PK4-1 | *W3110 ftsL::kan/pKTP108* | P1 on BL156 X W3110/pKTP108 |
| S3 | W3110 *leu*::Tn*10* | (6) |
| SD247 | W3110 *ftsW*$^{M269I}$ | (7) |
| SD247-1 | W3110 *recA::spc ftsW*$^{M269I}$ | P1 on KTP1 X SD247 select SpcR |
| SD285 | W3110 *leu*::Tn*10 bla lacI*$^{q}$ *P*$_{207}$-*gfp-ftsI* | P1 on EC436 X S3 (select AmpR) |
| SD399 | *W3110, ftsL::kan/*pSD256 | (7) |
| SD439 | W3110*, ftsL::kan*/pSD296 | (7) |

| | | |
|---|---|---|
| SD488 | S3 *ftsW*$^{E289G}$ | Integration and resolution of pSD257 in S3, screen small cell phenotype and Spc$^{S}$ |
| W3110 | WT | Lab collection |

B. Plasmids

| Plasmid | Genotype | Origin | Source/reference |
|---|---|---|---|
| pBAD33 | *cat* $P_{ara}$ | p15A | (8) |
| pBL154 | *cat* pSC101 *repA*$^{TS}$ $P_{syn135}$::*ftsN* | | (1) |
| pDSW208 | *bla lacI*$^{q}$ $P_{trc/204}$::*gfp* | ColE1 | (3) |
| pDSW210 | *bla lacI*$^{q}$ $P_{trc/206}$::*gfp* | ColE1 | (3) |
| pGB2 | *aadA* $P_{syn135}$::*vector* | pSC101 | (9) |
| pJF118EH | *bla lacI*$^{q}$ $P_{lac}$::*vector* | ColE1 | (10) |
| pKTP100 | pJF118EH $P_{tac}$::*ftsL* | ColE1 | This study |
| pKTP100* | pJF118EH $P_{tac}$:: *ftsL*$^{L86F,E87K}$ | ColE1 | This study |
| pKTP101 | pJF118EH $P_{tac}$::*ftsB* | ColE1 | This study |
| pKTP102 | pJF118EH $P_{tac}$::*torA*$^{1-42}$ *ftsL*$^{58-121}$-*6xhis* | ColE1 | This study |
| pKTP103 | pJF118EH $P_{tac}$::*malF*$^{1-37}$ *ftsL*$^{58-121}$-*6xhis* | ColE1 | This study |
| pQE80L | Expression vector with $P_{trc}$, Amp$^{R}$ | Qiagen | Qiagen |
| pKTP104 | *bla* pQE80L $P_{T5}$::*ftsL* | | This study |
| pKTP105 | *bla* pQE80L $P_{T5}$::*ftsL*$^{30-121}$ | | This study |
| pKTP106 | pBAD33 $P_{ara}$:: *ftsL* | p15A | This study |
| pKTP107 | pBAD33 $P_{ara}$:: *ftsL*$^{30-121}$ | p15A | This study |
| pKTP107X | pBAD33 $P_{ara}$::*ftsL*$^{\Delta 1-30}$-*6Xhis* | P15A | This study |
| pKTP108 | *aadA* pSC101 repA$^{TS}$ $P_{syn135}$:: *ftsL* | pSC101 | This study |
| pKTP109 | *cat* pBAD33 $P_{ara}$:: *ftsI* | P15A | This study |
| pND16 | *aadA* $P_{ftsK}$:: *ftsW-ftsK*$^{179-1329}$ | pSC101 | (11) |
| pND16* | *aadA* $P_{ftsK}$:: *ftsW*$^{M269I}$-*ftsK*$^{179-1329}$ | pSC101 | This study |
| pSD256 | *aadA repA*$^{ts}$ $P_{ftsL}$:: *ftsL* | pSC101 | (7) |

| | | | |
|---|---|---|---|
| pSD257 | *aadA repA$^{ts}$* P$_{ftsW}$*:: ftsW* | pSC101 | This study |
| pSD257* | *aadA repA$^{ts}$* P$_{ftsW}$*:: ftsW$^{M269I}$* | pSC101 | This study |
| pSD296 | *cat* pBAD33 P$_{ara}$*:: ftsL* | p15A | This study |
| pSD296-2 | *cat* pBAD33 P$_{ara}$*:: ftsL$^{L86F,E87K}$* | p15A | This study |
| pSEB417 | pDSW208 P$_{204}$*::ftsN* | ColE1 | (12) |
| pSEB420 | pDSW208 P$_{204}$*::ftsI* | ColE1 | (12) |
| pSEB422 | pDSW208 P$_{204}$*::ftsL* | ColE1 | (12) |
| pSEB429 | pDSW208 P$_{204}$:: *ftsW* | ColE1 | (12) |
| pSEB429-I | pDSW208 P$_{204}$:: *ftsW$^{M269I}$* | ColE1 | This study |
| pSEB453 | *bla* P$_{ftsN}$*:: malG$^{1–33}$–ftsN$^{46–319}$* | ColE1 | This study |
| pSEB468 | *aadA$^{s}$* P$_{syn135}$*:: ftsQ* | pSC101 | This work |
| pUT18 | *bla* BTH vector | ColE1 | (2) |
| pKT25C | *aph* BTH vector | p15A | (2) |
| pUT18-zip and pKT25-zip | BTH control plasmids | | (2) |
| pKT25C-ftsW | *aph* P$_{lac}$*::ftsW* | p15A | This work |
| pKT25C-ftsW* | *aph* P$_{lac}$*::ftsW$^{M269I}$* | p15A | This work |
| pKT25C-ftsI | *aph* P$_{lac}$*::ftsI* | P15A | This work |
| pUT18-ftsL | *bla* P$_{lac}$::*ftsL* | ColE1 | This work |
| pUT18-ftsL$^{\Delta 1-30}$ | *bla* P$_{lac}$::*ftsL$^{\Delta 1-30}$* | ColE1 | This work |

Table S2. List of the primers used in this study

| Primer Name | Sequence |
|---|---|
| *ftsL-Bam*HI-F | 5’-CGCGGGATCCATCAGCAGAGTGACAGAAGCTCTAA-3’ |
| *ftsL*$^{30-121}$-*Bam*HI-F | 5’-CGCG GGATCCGACGATCTTTTGCGATTTGGGAA-3’ |
| *ftsL*$^{30-121}$-*Bam*HI-F (B2H) | 5’-CGCGGGATCCAGACGATCTTTTGCGATTTGGGAA-3’ |
| *ftsL*$^{30-121}$-*Sac*I-F | 5’-CGCG GAGCTC ATTGCAGAGAGGACGAATGC ATG GAC GATCTTTTGCGATTTGGGAA-3’ |
| *ftsL-Eco*RI-F | 5’-CGCGGAATTC ATTGCAGAGAGGACGAATGCATGATCAGCAGAGT GACAGA-3’ |
| *ftsL-Hin*dIII-R | 5’-CAGTAAGCTT CTA TTTTTGCACTACGATATTTTCTT-3’ |
| *ftsL-Hin*dIII-F | 5’-CTTTACTAAGCTTCGTATTGTGAAACGTTTTATGCGT-3’ |
| *ftsL-Eco*RI-R | 5’-GTCGGGAATTCAGAGAACGCATGTCGCCCTCTT-3’ |
| *ftsL-Sac*I-F | 5’-CGCGGAGCTC ATTAAAGAGGAGAAATTAACT ATGATCAGCAGAGTG ACAGA-3’ |
| *ftsL-Xba*I-F | 5’-GATCCTCTAGAATTGCAGAGAGGACGAATGCATGA-3’ |
| *ftsL-Hin*dIII-R (pSD296) | 5’-CCAAGCTTATTTTTGCACTACGATATTTT-3’ |
| *malF*$^{1-37}$*ftsL*$^{58-121}$-*Eco*RI-F | 5’- CGCGGAATTCATTGCAGAGAGGACGAATGCATGGATGTCATTAAA AAGAAACATTGGTGGCAAAGCGACGCGCTGAAATGGTCAGTGCTAGG TCTGCTCGGCCTGCTGGTGGGTTACCTTGTTGTTTTAATGTACGCACAA CACCATACCCGTTTACTGACCGCTCAGCGCGAACAACTG-3’ |
| *torA*$^{1-42}$ *ftsL*$^{58-12L}$-*Eco*RI- F | 5’- CGCGGAATTC ATTGCAGAGAGGACGAATGCATGAACAATAACGAT CTCTTTCAGGCATCACGTCGGCGTTTTCTGGCACAACTCGGCGGCTT AACCGTCGCCGGGATGCTGGGGCCGTCATTGTTAACGCCGCGACGT GCGACTGCGGCGCAAGCGCACCATACCCGTTTACTGACCGCTCAGC GCGAACAACTG-3’ |
| *ftsL-6xhis-Hin*dIII-R | 5’- CAGTAAGCTTCTAGTGATGGTGATGGTGATGTTTTTGCACTACGA TATTTTCTT-3’ |
| *ftsI-Sac*I-F | CGCGGAGCTCATTAAAGAGGAGAAATTAACTATGAAAGCAGCGGCGAAA A |
| *ftsI-Hind*III-R | CGCGAAGCTTTCACGATCTGCCACCTGTCCCCTCGCC |
| *ftsW-Bam*HI-F | 5’-CGCGGGATCCACGTTTATCTCTCCCTCGCCTGAAAATG-3’ |
| *ftsW-Kpn*I-R | 5’-CGCGGGTACCTCATCGTGAACCTCGTACAAACGCCTGC-3’ |

| | |
|---|---|
| *ftsI-Bam*HI-F | 5’-CGCGGGATCCAAAAGCAGCGGCGAAAACGCAG-3’ |
| *ftsI-Eco*RI-R | 5’-CGCGGAATTCTCACGATCTGCCACCTGTCCCCTCGCC-3’ |
| *ftsB-BamH*I-F | 5’-CGCGGGATCCGGTAAACTAACGCTGCTGTTGCTG-3’ |
| *ftsB-Eco*RI-F | 5’-CGCGGAATTCGCCGTTTTTCAGGGGGCAGGATGGGTAAAC-3’ |
| *ftsB-Hin*dIII-R | 5’-CAGTAAGCTTTTATCGATTGTTTTGCCCCGCAGACTGTGCGC-3’ |
| *ftsB*-N43K-F | 5’-GCACAGCAAGCTACAAAAGCGAAACTTAAAGCG-3’ |
| *ftsB*-N43K-R | 5’-GCTTTAAGTTTCGCTTTTGTAGCTTGCTGTGCC-3’ |
| *ftsB*-N50K-F | 5’-AACTTAAAGCGCGAAAAGATCAACTTTTTGCCG-3’ |
| *ftsB*-N50K-R | 5’-CGGCAAAAAGTTGATCTTTTCGCGCTTTAAGTT-3’ |
| *ftsB*-Q52K-F | 5’-GCGAAACGATAAACTTTTTGCCGAAATTGACGA-3’ |
| *ftsB*-Q52K-R | 5’-TCGTCAATTTCGGCAAAAAGTTTATCGTTTCGC-3’ |
| *ftsB*-F54K-F | 5’-AAACGATCAACTTAAAGCCGAAATTGACGATCTC-3’ |
| *ftsB*-F54K-R | 5’-GAGATCGTCAATTTCGGCTTTAAGTTGATCGTTT-3’ |
| *ftsB*-E56A-F | 5’-ATCAACTTTTTGCCGCAATTGACGATCTCAATG-3’ |
| *ftsB*-E56A-R | 5’-CATTGAGATCGTCAATTGCGGCAAAAAGTTGAT-3’ |
| *ftsB*-I57K-F | 5’-AACTTTTTGCCGAAAAAGACGATCTCAATGGCG-3’ |
| *ftsB*-I57K-R | 5’-CGCCATTGAGATCGTCTTTTTCGGCAAAAAGTT-3’ |
| *ftsB*-L60K-F | 5’-CGAAATTGACGATAAAAATGGCGGCCAGGAGGC-3’ |
| *ftsB*-L60K-R | 5’-GCCTCCTGGCCGCCATTTTTATCGTCAATTTCG-3’ |

| | |
|---|---|
| *ftsB*-A66K-F | 5’-TCAATGGCGGCCAGGAGAAGCTCGAAGAGCGTGC-3’ |
| *ftsB*-A66K-R | 5’-GCACGCTCTTCGAGCTTCTCCTGGCCGCCATTGA-3’ |
| *ftsB*-R72A-F | 5’-GAAGAGCGTGCGGCTAATGAACTCAGCATGACC-3’ |
| *ftsB*-R72A-R | 5’-GGTCATGCTGAGTTCATTAGCCGCACGCTCTTC-3’ |
| *ftsB*-E82A-F | 5’-TGACCAGGCCGGGCGCAACTTTTTATCGTC-3’ |
| *ftsB*-E82A-R | 5’-GACGATAAAAAGTTGCGCCCGGCCTGGTCA-3’ |
| *ftsL*-I28E-F | 5’-GCATTGCCTGGTGTTGAAGGTGACGATCTTTTG-3’ |
| *ftsL*-I28E-R | 5’-CAAAAGATCGTCACCTTCAACACCAGGCAATGC-3’ |
| *ftsL*-I28K-F | 5’-GCATTGCCTGGTGTTAAAGGTGACGATCTTTTG-3’ |
| *ftsL*-I28K-R | 5’-CAAAAGATCGTCACCTTTAACACCAGGCAATGC-3’ |
| *ftsL*-L24K/I28K-F | 5’-CGCCATGCAAAGCCTGGTGTTAAAGGTGACGATCTT-3’ |
| *ftsL*-L24K/I28K-R | 5’-AAGATCGTCACCTTTAACACCAGGCTTTGCATGGCG-3’ |
| *ftsL*-R61E-F | 5’-GTAACCACGGCGCACCATACCGAATTACTGACCGC-3’ |
| *ftsL*-R61E-R | 5’-GCGGTCAGTAATTCGGTATGGTGCGCCGTGGTTAC-3’ |
| *ftsL*-L77K-F | 5’-GCTGGAGCGAGATGCTAAAGACATTGAATG-3’ |
| *ftsL*-L77K-R | 5’-CATTCAATGTCTTTAGCATCTCGCTCCAGC-3’ |
| *ftsL*-D78K-F | 5’-GAGCGAGATGCTTTAAAAATTGAATGGCGCA-3’ |
| *ftsL*-D78K-R | 5’-TGCGCCATTCAATTTTTAAAGCATCTCGCTC-3’ |

| | |
|---|---|
| *ftsL*-E80K-F | 5’-ATGCTTTAGACATTAAATGGCGCAACCTGA-3’ |
| *ftsL*-E80K-R | 5’-TCAGGTTGCGCCATTTAATGTCTAAAGCAT-3’ |
| *ftsL*-W81A-F | 5’-CTTTAGACATTGAAGCGCGCAACCTGATCCTT-3’ |
| *ftsL*-W81A-R | 5’-AAGGATCAGGTTGCGCGCTTCAATGTCTAAAG-3’ |
| *ftsL*-R82E-F | 5’-GACATTGAATGGGAAAACCTGATCCTTGAAGAG-3’ |
| *ftsL*-R82E-R | 5’-CTCTTCAAGGATCAGGTTTTCCCATTCAATGTC-3’ |
| *ftsL*-N83K-F | 5’-ATTGAATGGCGCAAACTGATCCTTGAAGAGAAT-3’ |
| *ftsL*-N83K-R | 5’-ATTCTCTTCAAGGATCAGTTTGCGCCATTCAAT-3’ |
| *ftsL*-N83K/E88K-F | 5’-TGAATGGCGCAAACTGATCCTTGAAAAGAATGCGCT-3’ |
| *ftsL*-N83K/E88K-R | 5’-AGCGCATTCTTTTCAAGGATCAGTTTGCGCCATTCA-3’ |
| *ftsL*-L84K-F | 5’-ATTGAATGGCGCAACAAGATCCTTGAAGAGA-3’ |
| *ftsL*-L84K-R | 5’-TCTCTTCAAGGATCTTGTTGCGCCATTCAAT-3’ |
| *ftsL*-L86F-F | 5’-TGGCGCAACCTGATCTTTGAAGAGAATGCGCT-3’ |
| *ftsL*-L86F-R | 5’-AGCGCATTCTCTTCAAAGATCAGGTTGCGCCA-3’ |
| *ftsL*-L86F/E88K-F | 5’-TGAATGGCGCAACCTGATCTTTGAAAAGAATGCG-3’ |
| *ftsL*-L86F/E88K-R | 5’-CGCATTCTTTTCAAAGATCAGGTTGCGCCATTCA-3’ |
| *ftsL*-L86F/G92D-F | 5’-ACCTGATCTTTGAAGAGAATGCGCTCGACGACCATAG-3’ |
| *ftsL*-L86F/G92D-R | 5’-CTATGGTCGTCGAGCGCATTCTCTTCAAAGATCAGGT-3’ |

| | |
|---|---|
| *ftsL*-E87A-F | 5’-ACCTGATCCTTGCAGAGAATGCGCTCGGCGACCAT-3’ |
| *ftsL*-E87A-R | 5’-ATGGTCGCCGAGCGCATTCTCTGCAAGGATCAGGT-3’ |
| *ftsL*-E87D-F | 5’-CCTGATCCTTGATGAGAATGCGCTCGGCGACCAT-3’ |
| *ftsL*-E87D-R | 5’-ATGGTCGCCGAGCGCATTCTCATCAAGGATCAGG-3’ |
| *ftsL*-E87K-F | 5’-CTGATCCTTAAAGAGAATGCGCTCGGCGACCAT-3’ |
| *ftsL*-E87K-R | 5’-ATGGTCGCCGAGCGCATTCTCTTTAAGGATCAG-3’ |
| *ftsL*-E87S-F | 5’-CTGATCCTTTCAGAGAATGCGCTCGGCGACCAT-3’ |
| *ftsL*-E87S-R | 5’-ATGGTCGCCGAGCGCATTCTCTGAAAGGATCAG-3’ |
| *ftsL*-E87T-F | 5’-ACCTGATCCTTACAGAGAATGCGCTCGGCGACCAT-3’ |
| *ftsL*-E87T-R | 5’-ATGGTCGCCGAGCGCATTCTCTGTAAGGATCAGGT-3’ |
| *ftsL*-E87K/L86F-F | 5’-GCAACCTGATCTTTAAAGAGAATGCGCTCGGCGACCAT-3’ |
| *ftsL*-E87K/L86F-R | 5’-ATGGTCGCCGAGCGCATTCTCTTTAAAGATCAGGTTGC-3’ |
| *ftsL*-E87K/E88K-F | 5’-GCAACCTGATCCTTAAAAAGAATGCGCTCGGCGACCAT-3’ |
| *ftsL*-E87K/E88K-R | 5’-ATGGTCGCCGAGCGCATTCTTTTTAAGGATCAGGTTGC-3’ |
| *ftsL*-L86F/E87K/E88K-F | 5’-AACCTGATCTTTAAAAAGAATGCGCTCGGCGACCAT-3’ |
| *ftsL*-L86F/E87K/E88K-R | 5’-ATGGTCGCCGAGCGCATTCTTTTTAAAGATCAGGTT-3’ |
| *ftsL*-E87K/G92A-F | 5’-ATCCTTAAAGAGAATGCGCTCGCCGACCATAGCCGG-3’ |
| *ftsL*-E87K/G92A-R | 5’-CCGGCTATGGTCGGCGAGCGCATTCTCTTTAAGGAT-3’ |

| | |
|---|---|
| *ftsL*-E87K/G92D-F | 5’-ATCCTTAAAGAGAATGCGCTCGACGACCATAGCCGG-3’ |
| *ftsL*-E87K/G92D-R | 5’-CCGGCTATGGTCGTCGAGCGCATTCTCTTTAAGGAT-3’ |
| *ftsL*-E87K/E88K/G92D-F | 5’-CTGATCCTTAAAAAGAATGCGCTCGACGACCAT-3’ |
| *ftsL*-E87K/E88K/G92D-R | 5’-ATGGTCGTCGAGCGCATTCTTTTTAAGGATCAG-3’ |
| *ftsL*-L86F/E87K/E88KG92D -F | 5’-AACCTGATCTTTAAAAAGAATGCGCTCGACGACCATAG-3’ |
| *ftsL*-L86F/E87K/E88KG92D -R | 5’-CTATGGTCGTCGAGCGCATTCTTTTTAAAGATCAGGTT-3’ |
| *ftsL*-E87K/G92E-F | 5’-ATCCTTAAAGAGAATGCGCTCGAAGACCATAGCCGG-3’ |
| *ftsL*-E87K/G92E-R | 5’-CCGGCTATGGTCTTCGAGCGCATTCTCTTTAAGGAT-3’ |
| *ftsL*-E87K/G92K-F | 5’-ACCTGATCCTTAAAGAGAATGCGCTCAAAGACCAT-3’ |
| *ftsL*-E87K/G92K-R | 5’-ATGGTCTTTGAGCGCATTCTCTTTAAGGATCAGGT-3’ |
| *ftsL*-E87K/G92S-F | 5’-ACCTGATCCTTAAAGAGAATGCGCTCAGCGACCAT-3’ |
| *ftsL*-E87K/G92S-R | 5’-ATGGTCGCTGAGCGCATTCTCTTTAAGGATCAGGT-3’ |
| *ftsL*-E87K/G92V-F | 5’-CCTGATCCTTAAAGAGAATGCGCTCGTCGACCATAGC-3’ |
| *ftsL*-E87K/G92V-R | 5’-GCTATGGTCGACGAGCGCATTCTCTTTAAGGATCAGG-3’ |
| *ftsL*-E88K-F | 5’-ACCTGATCCTTGAAAAGAATGCGCTCGGCGACCAT-3’ |
| *ftsL*-E88K-R | 5’-ATGGTCGCCGAGCGCATTCTTTTCAAGGATCAGGT-3’ |
| *ftsL*-E88K/A90E-F | 5’-TGATCCTTGAAAAGAATGAGCTCGGCGACCATA-3’ |
| *ftsL*-E88K/A90E-R | 5’-TATGGTCGCCGAGCTCATTCTTTTCAAGGATCA-3’ |

| | |
|---|---|
| *ftsL*-A90E-F | 5’-CTTGAAGAGAATGAGCTCGGCGACCATAGCCGG-3’ |
| *ftsL*-A90E-R | 5’-CCGGCTATGGTCGCCGAGCTCATTCTCTTCAAG-3’ |
| *ftsL*-A90G-F | 5’-TCCTTGAAGAGAATGGGCTCGGCGACCATAGCC-3’ |
| *ftsL*-A90G-R | 5’-GGCTATGGTCGCCGAGCCCATTCTCTTCAAGGA-3’ |
| *ftsL*-A90I-F | 5’-TCCTTGAAGAGAATATTCTCGGCGACCATAGCC-3’ |
| *ftsL*-A90I-R | 5’-GGCTATGGTCGCCGAGAATATTCTCTTCAAGGA-3’ |
| *ftsL*-A90K-F | 5’-CTTGAAGAGAATAAGCTCGGCGACCATAGCCGG-3’ |
| *ftsL*-A90K-R | 5’-CCGGCTATGGTCGCCGAGCTTATTCTCTTCAAG-3’ |
| *ftsL*-A90L-F | 5’-CCTTGAAGAGAATCTGCTCGGCGACCATAGCCG-3’ |
| *ftsL*-A90L-R | 5’-CGGCTATGGTCGCCGAGCAGATTCTCTTCAAGG-3’ |
| *ftsL*-L91K-F | 5’-TTGAAGAGAATGCGAAAGGCGACCATAGCCG-3’ |
| *ftsL*-L91K-R | 5’-CGGCTATGGTCGCCTTTCGCATTCTCTTCAA-3’ |
| *ftsL*-G92D-F | 5’-CTGATCCTTGAAGAGAATGCGCTCGACGACCATAG-3’ |
| *ftsL*-G92D-R | 5’-CTATGGTCGTCGAGCGCATTCTCTTCAAGGATCAG-3’ |
| *ftsL*-E88K/G92D-F | 5’-CTGATCCTTGAAAAGAATGCGCTCGACGACCATAG-3’ |
| *ftsL*-E88K/G92D-R | 5’-CTATGGTCGTCGAGCGCATTCTTTTCAAGGATCAG-3’ |
| *ftsL*-A90E/G92D-F | 5’-CTTGAAGAGAATGAGCTCGACGACCATAGCCGG-3’ |
| *ftsL*-A90E/G92D-R | 5’-CCGGCTATGGTCGTCGAGCTCATTCTCTTCAAG-3’ |

| | |
|---|---|
| *ftsL*-A90E/E88K/G92D-F | 5’-ACCTGATCCTTGAAAAGAATGAGCTCGACGACCATAG-3’ |
| *ftsL*-A90E/E88K/G92D-R | 5’-CTATGGTCGTCGAGCTCATTCTTTTCAAGGATCAGGT-3’ |
| *ftsL*-G92R-F | 5’-TGAAGAGAATGCGCTCCGCGACCATAGCCGGGTG-3’ |
| *ftsL*-G92R-R | 5’-CACCCGGCTATGGTCGCGGAGCGCATTCTCTTCA-3’ |
| *ftsL*-D93K-F | 5’-TGAAGAGAATGCGCTCGGCAAACATAGCCGGGTG-3’ |
| *ftsL*-D93K-R | 5’-CACCCGGCTATGTTTGCCGAGCGCATTCTCTTCA-3’ |
| *ftsL*-R96E-F | 5’-CGACCATAGCGAGGTGGAAAGGATCGCCACGGA-3’ |
| *ftsL*-R96E-R | 5’-TCCGTGGCGATCCTTTCCACCTCGCTATGGTCG-3’ |
| *ftsL*-I100 stop-F | 5’-CCATAGCCGGGTGGAAAGGTAGTAGACGGAAAAG-3’ |
| *ftsL*-I100 stop-R | 5’-CTTTTCCGTCTACTACCTTTCCACCCGGCTATGG-3’ |
| *ftsL*-A101K-F | 5’-GGGTGGAAAGGATCAAAACGGAAAAGCTGCAAATGCA-3’ |
| *ftsL*-A101K-R | 5’-TGCATTTGCAGCTTTTCCGTTTTGATCCTTTCCACCC-3’ |
| *ftsL*-L105D-F | 5’-TCGCCACGGAAAAGGACCAAATGCAGCATGTTG-3’ |
| *ftsL*-L105D-R | 5’-CAACATGCTGCATTTGGTCCTTTTCCGTGGCGATC-3’ |
| *ftsL*-M107A-F | 5’-CCACGGAAAAGCTGCAAGCGCAGCATGTTGATC-3’ |
| *ftsL*-M107A-R | 5’-GATCAACATGCTGCGCTTGCAGCTTTTCCGTGG-3’ |
| *ftsL*-M107D-F | 5’-CCACGGAAAAGCTGCAAGACCAGCATGTTGATC-3’ |
| *ftsL*-M107D-R | 5’-GATCAACATGCTGGTCTTGCAGCTTTTCCGTGG-3’ |

| | |
|---|---|
| *ftsL*-M107E-F | 5’-CCACGGAAAAGCTGCAAGAGCAGCATGTTGATC-3’ |
| *ftsL*-M107E-R | 5’-ATCAACATGCTGCTCTTGCAGCTTTTCCGTGG-3’ |
| *ftsL*-M107G-F | 5’-CCACGGAAAAGCTGCAAGGGCAGCATGTTGATC-3’ |
| *ftsL*-M107G-R | G5’-ATCAACATGCTGCCCTTGCAGCTTTTCCGTGG-3’ |
| *ftsL*-M107K-F | 5’-CCACGGAAAAGCTGCAAAAGCAGCATGTTGAT-3’ |
| *ftsL*-M107K-R | 5’-ATCAACATGCTGCTTTTGCAGCTTTTCCGTGG-3’ |
| *ftsL*-M107R-F | 5’-CCACGGAAAAGCTGCAACGGCAGCATGTTGATC-3’ |
| *ftsL*-M107R-R | 5’-GATCAACATGCTGCCGTTGCAGCTTTTCCGTGG-3’ |
| *ftsL*-M107 stop-F | 5’-CCACGGAAAAGCTGCAATAGCAGCATGTTGAT-3’ |
| *ftsL*-M107 stop-R | 5’-ATCAACATGCTGCTATTGCAGCTTTTCCGTGG-3’ |
| *ftsL*-E115G-F | 5’-GTTGATCCGTCACAAGGAAATATCGTAGTGCAA-3’ |
| *ftsL*-E115G-R | 5’-TTGCACTACGATATTTCCTTGTGACGGATCAAC-3’ |
| *ftsL*-E115K-F | 5’-GTTGATCCGTCACAAAAAAATATCGTAGTGCAA-3’ |
| *ftsL*-E115K-R | 5’-TTGCACTACGATATTTTTTTGTGACGGATCAAC-3’ |
| *ftsL*-E115R-F | 5’-GTTGATCCGTCACAACGAAATATCGTAGTGCAA-3’ |
| *ftsL*-E115R-R | 5’-TTGCACTACGATATTTCGTTGTGACGGATCAAC-3’ |
| *ftsL*-P112 stop-F | 5’-TGCAGCATGTTGATTAGTCACAAGAAAATATCG-3’ |
| *ftsL*-P112 stop-R | 5’-CGATATTTTCTTGTGACTAATCAACATGCTGCA-3’ |

| | |
|---|---|
| *ftsL*-Q114 stop-F | 5'-GCATGTTGATCCGTCATAGGAAAATATCGTAGT-3' |
| *ftsL*-Q114 stop-R | 5'-ACTACGATATTTTCCTATGACGGATCAACATGC-3' |
| *ftsW*-M269I-F | 5'-CGCAATCGCTGATCGCGTTTGGTCGCGGCGAACTT-3' |
| *ftsW*-M269I-R | 5'-AAGTTCGCCGCGACCAAACGCGATCAGCGATTGCG-3' |
| *ftsW*-E289G-F | 5'-TCGGTACAAAAACTGGGGTATCTGCCGGAAGCG-3' |
| *ftsW*-E289G-R | 5'-CGCTTCCGGCAGATACCCCAGTTTTTGTACCGA-3' |
| *ftsW*-E289K-F | 5'-TCGGTACAAAAACTGAAGTATCTGCCGGAAGCG-3' |
| *ftsW*-E289K-R | 5'-CGCTTCCGGCAGATACTTCAGTTTTTGTACCGA-3' |
| *ftsW*-E289R-F | 5'-TCGGTACAAAAACTGCGGTATCTGCCGGAAGCG-3' |
| *ftsW*-E289R-R | 6'-CTTCCGGCAGATACCGCAGTTTTTGTACCGA-3' |

1. Gerding MA*, et al.* (2009) Self-enhanced accumulation of FtsN at Division Sites and Roles for Other Proteins with a SPOR domain (DamX, DedD, and RlpA) in Escherichia coli cell constriction. *J Bacteriol* 191(24):7383-7401.
2. Karimova G, Dautin N, & Ladant D (2005) Interaction network among Escherichia coli membrane proteins involved in cell division as revealed by bacterial two-hybrid analysis. *J Bacteriol* 187(7):2233-2243.
3. Weiss DS, Chen JC, Ghigo JM, Boyd D, & Beckwith J (1999) Localization of FtsI (PBP3) to the septal ring requires its membrane anchor, the Z ring, FtsA, FtsQ, and FtsL. *J Bacteriol* 181(2):508-520.
4. Pichoff S & Lutkenhaus J (2007) Identification of a region of FtsA required for interaction with FtsZ. *Mol Microbiol* 64(4):1129-1138.
5. Dai K, Xu Y, & Lutkenhaus J (1993) Cloning and characterization of ftsN, an essential cell division gene in Escherichia coli isolated as a multicopy suppressor of ftsA12(Ts). *J Bacteriol* 175(12):3790-3797.

6. Shen B & Lutkenhaus J (2009) The conserved C-terminal tail of FtsZ is required for the septal localization and division inhibitory activity of MinC(C)/MinD. *Mol Microbiol* 72(2):410-424.
7. Du S, Pichoff S, & Lutkenhaus J (2016) FtsEX acts on FtsA to regulate divisome assembly and activity. *Proc Natl Acad Sci U S A* 113(34):E5052-5061.
8. Guzman LM, Belin D, Carson MJ, & Beckwith J (1995) Tight regulation, modulation, and high-level expression by vectors containing the arabinose PBAD promoter. *J Bacteriol* 177(14):4121-4130.
9. Churchward G, Belin D, & Nagamine Y (1984) A pSC101-derived plasmid which shows no sequence homology to other commonly used cloning vectors. *Gene* 31(1-3):165-171.
10. Furste JP*, et al.* (1986) Molecular cloning of the plasmid RP4 primase region in a multi-host-range tacP expression vector. *Gene* 48(1):119-131.
11. Dubarry N, Possoz C, & Barre FX (2010) Multiple regions along the Escherichia coli FtsK protein are implicated in cell division. *Mol Microbiol* 78(5):1088-1100.
12. Pichoff S, Du S, & Lutkenhaus J (2015) The bypass of ZipA by overexpression of FtsN requires a previously unknown conserved FtsN motif essential for FtsA-FtsN interaction supporting a model in which FtsA monomers recruit late cell division proteins to the Z ring. *Mol Microbiol* 95(6):971-987.
